# LKB1 Spatially Regulates AMPK Activity Coordinate Cellular Oxidative Stress Response

**DOI:** 10.1101/2025.11.17.688926

**Authors:** Kasey Parks, Arnav Jhawar, Alexia Andrikopoulos, Declan M. Winters, Teagan S. Dean, Edmund D. Kapelczak, Tara TeSlaa, Mehdi Bouhaddou, Danielle L. Schmitt

## Abstract

AMP-activated protein kinase (AMPK) is a central regulator of cellular energy homeostasis, with over 100 identified downstream targets throughout the cell. In response to cellular stress, including energetic stress, AMPK is activated via binding of AMP and phosphorylation by upstream kinases, including liver kinase B1 (LKB1). We and others have found that the activation of AMPK in response to cellular stress has distinct subcellular mechanisms, indicating compartmentalized regulation of AMPK signaling. Oxidative stress is known to stimulate AMPK activity, but how AMPK is spatially regulated by oxidative stress is underexplored. Using a single-fluorophore excitation-ratiometric AMPK activity reporter (ExRai AMPKAR), we find that oxidative stress induced by hydrogen peroxide (H_2_O_2_) results in AMPK activity with distinct spatiotemporal dynamics. We found that in the cytoplasm, nucleus, outer mitochondrial membrane, and cytosolic lysosomal surface, phosphorylation of AMPK by LKB1 is required for AMPK activity. Using a biosensor for ATP, we found at the cytoplasm and lysosome local ATP depletion dictates kinetics of AMPK activity. Using a multi-‘omics approach, we discover that in response to oxidative stress, AMPK mediates significant metabolic and gene expression changes, including upregulation of oxidative stress response through nuclear factor erythroid 2-related factor 2 (NRF2). Expanding on this identified mechanism, we find that non-small cell lung cancers harboring Kelch-like ECH-associated protein 1 (KEAP1) mutations have a functionally deficient LKB1-AMPK signaling network in response to oxidative stress. Altogether, this work provides new insights into how the subcellular environment influences localized AMPK activity, and identifies how AMPK regulates the cellular response to oxidative stress.

## Introduction

AMP-activated protein kinase (AMPK) is a master regulator of cellular energy homeostasis, coordinating metabolic adaptation to nutrient deprivation, mitochondrial dysfunction, and stress.^1,2^ An obligate heterotrimeric protein, AMPK consists of an α catalytic subunit, β regulatory subunit, and γ nucleotide binding subunit. In mammals, these subunits exist in multiple isoforms (α1/α2, β1/β2, γ1/γ2/γ3), leading to at least 12 distinct AMPK complexes.^3–5^ Under low-energy conditions, rising AMP levels promote AMPK activation through binding of AMP to the γ subunit and protection from dephosphorylation of Thr172 on the ɑ subunit, which is canonically phosphorylated by the upstream kinase liver kinase B1 (LKB1) and calcium/calmodulin-dependent protein kinase kinase 2 (CaMKK2).^6–9^ CaMKK2 can also activate AMPK through AMP-independent mechanisms.^10^ Once active, AMPK restores energy homeostasis by modulating activity of metabolic enzymes, mitochondrial fision and fusion, transcriptional programs, and autophagy.^1^

Regulation of cellular energy homeostasis by AMPK is linked to its tight spatiotemporal regulation within distinct subcellular compartments.^11,12^ To explore compartmentalized AMPK activity, genetically encoded AMPK activity reporters (AMPKARs) allow for real time visualization of AMPK activity in single cells, with subcellular resolution achievable using localization sequences for cellular organelles and compartments.^13^ More recently, we reported on the development of an excitation-ratiometric AMPK activity reporter (ExRai AMPKAR), which we used identify distinct mechanisms for AMPK activity based on location and upstream kinase in response to energy stress or allosteric activation.^14^ For instance, we found that LKB1 mediated rapid AMPK activity at the lysosome, as loss of LKB1 resulted in slower accumulation of AMPK activity, which likely represents a priveledged site for AMPK sensing of the cellular energetic state.^14^ Thus, measuring localized AMPK activity in response to different activating conditions is crucial for fully understanding the complex regulatory network of AMPK.

While the spatiotemporal regulation of AMPK in response to energy stress has been described, mechanisms for spatial AMPK activity under oxidative stress remain poorly defined. AMPK has been well documented to become active in response to oxidative stress, usually induced by hydrogen peroxide treatment. Some studies proposed ATP depletion under oxidative stress leads to AMPK activation via AMP/ATP imbalance.^15,16^ Others suggested mechanisms involving direct oxidation of cystine residues on AMPK or its upstream kinases.^9,11,17–19^ Additionally, some studies have indicated that LKB1 is important for oxidative stress-induced AMPK activity.^9,19^ However, how subcellular location influences AMPK activity in response to oxidative stress remains unclear.

Here, we investigate mechanisms for spatial AMPK activation by oxidative stress. Using subcellularly localized ExRai AMPKAR, we find that oxidative stress-induced AMPK activation has distinct spatial dynamics in the cytoplasm, nucleus, mitochondria, and lysosome. We find that oxidative stress-induced AMPK activity is dependent on phosphorylation by LKB1. We use a complimentary multi-‘omics approach to identify downstream consequences of oxidative stress-induced AMPK activity, including regulation of fatty acid biosynthesis, the pentose phosphate pathway, and NRF2-mediated gene expression. Finally, we find our identified mechanism for oxidative stress-induced AMPK activation is defective in KEAP1/LKB1/KRAS driven non-small cell lung cancers, illuminating how KEAP1 mutations influence AMPK signaling. Collectively, our results clarify the spatial activation of AMPK under oxidative stress and delineate how AMPK activation reprograms cell function in response to oxidative stress.

## Results

### AMPK has distinct subcellular activity dynamics in response to oxidative stress

To examine how oxidative stress influences AMPK activity at distinct cellular compartments in live cells, we used our excitation-ratiometric AMPK activity reporter (ExRai AMPKAR).^14^ With organelle-targeted ExRai AMPKAR, we assessed spatial AMPK activity in the cytoplasm, nucleus, mitochondrial outer membrane, and lysosomal surface of mouse embryonic fibroblasts (MEFs) following hydrogen peroxide (H_2_O_2_) exposure (Fig. 1a). Stimulation with H_2_O_2_ induced robust increases in the ExRai AMPKAR excitation-emission ratio (R = 480 nm ex-em/400 nm ex-em) and maximum ratio change (ΔR/R_0_) in all compartments tested (Fig. 1b-e).

**Figure 1.**
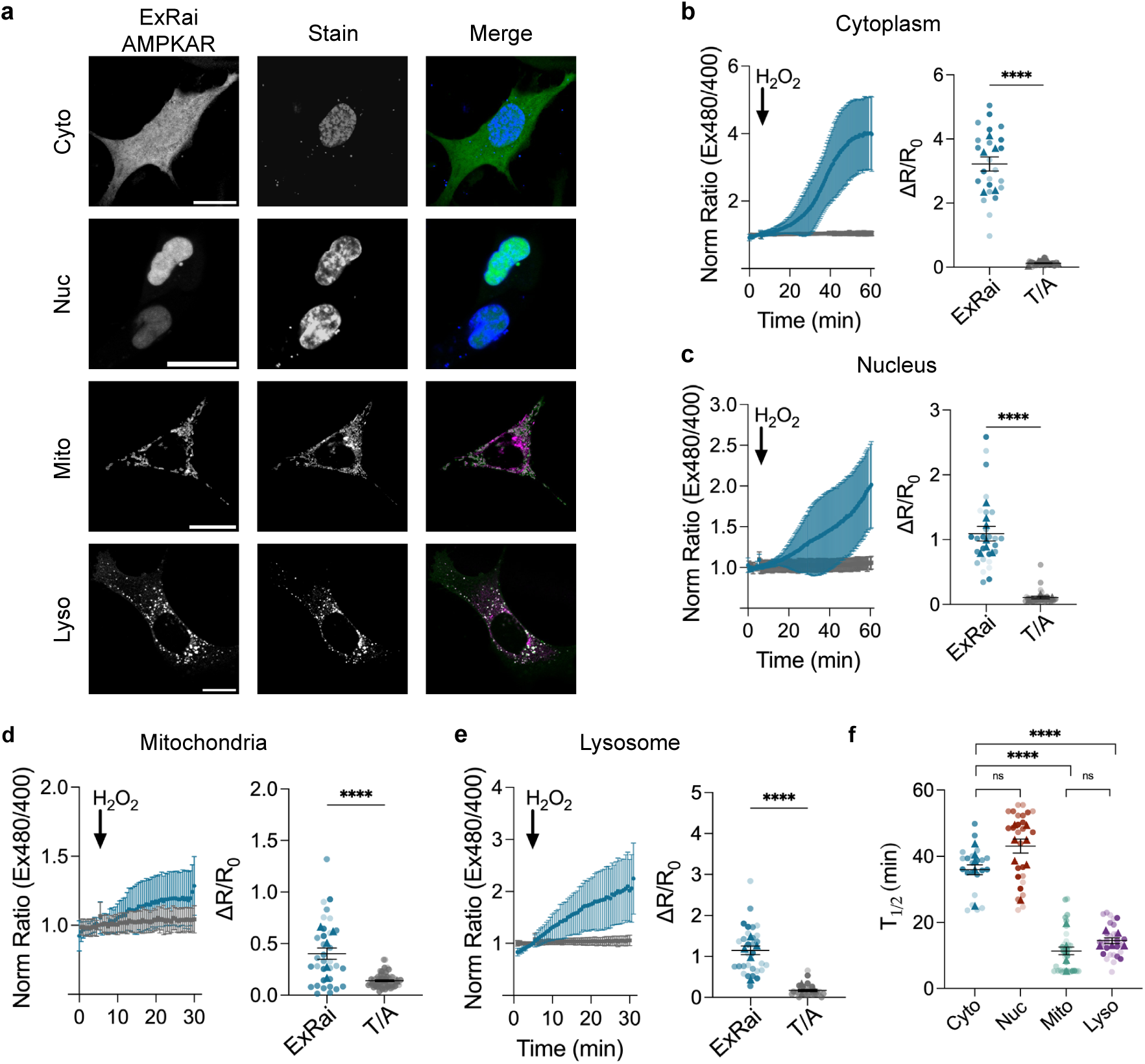
AMPK exhibits compartmentalized responses to oxidative stress. **a** (From top to bottom) Representative images of: ExRai AMPKAR in MEFs stained with a DAPI nuclear stain; nuclear-locatized ExRai AMPKAR (NLS-ExRai AMPKAR) in MEFs stained with a DAPI nuclear stain; outer mitochondrial membrane-localized ExRai AMPKAR (mito-ExRai AMPKAR) in MEFs stained with the mitochondrial marker MitoTracker Red; outer lysosomal membrane-localized ExRai AMPKAR (lyso-ExRai AMPKAR) stained with the lysosomal marker LysoTracker Red. Scale bars represent 20 µm. **b** Average response of ExRai AMPKAR (blue, n=23 cells from five experiments) and ExRai AMPKAR T/A (grey, n=39 cells from four experiments) in WT MEFs treated with H_2_O_2_ (150µM), along with maximum ratio change (\*\*\*\**p*<0.0001, unpaired *t*-test, two-tailed). **c** Average response of NLS-ExRai AMPKAR (blue, n=27 cells from seven experiments) and NLS-ExRai AMPKAR T/A (grey, n=32 cells from three experiments) in WT MEFs treated with H_2_O_2_ (150µM), along with maximum ratio change (\*\*\*\**p*<0.0001, unpaired *t*-test, two-tailed). **d** Average response of mito-ExRai AMPKAR (blue, n=33 cells from five experiments) and mito-ExRai AMPKAR T/A (grey, n=60 cells from three experiment) in WT MEFs treated with H_2_O_2_ (150µM), along with maximum ratio change (\*\**p*=0.0051, unpaired *t*-test, two-tailed). **e** Average response of lyso-ExRai AMPKAR (blue, n=30 cells from five experiments) and lyso-ExRai AMPKAR T/A (grey, n=31 cells from three experiments) in WT MEFs treated with H_2_O_2_ (150µM), along with maximum ratio change (\*\*\*\**p*<0.0001, unpaired *t*-test, two-tailed). **f** *t*_1/2_ of cytoplasmic ExRai AMPKAR (blue), NLS-ExRai AMPKAR (red), mito-ExRai AMPKAR (teal), and lyso-ExRai AMPKAR (purple) from b-e following treatment with 150µM H_2_O_2_ (ns≥0.1508, \*\*\*\**p*<0.0001, unpaired *t*-test, two-tailed). For all figures, time courses show the mean ± SD, dot plots show the mean ± SEM.

In contrast, cells expressing a phospho-null ExRai AMPKAR where the AMPK phosphorylation site T is mutated to A (T/A) exhibited negligible changes in ExRai AMPKAR response, confirming that the observed signals reflect AMPK activity. In comparing across compartments, we observed differences in AMPK activation kinetics. The time-to-half maximal response (*t*_1/2_) of ExRai AMPKAR was significantly faster at the lysosome (*t*_1/2_=14.48 ± 0.86 min) and mitochondria (*t*_1/2_=11.36 ± 1.14 min) compared to the nucleus (*t*_1/2_=41.67 ± 2.45 min) and cytoplasm (*t*_1/2_=36.52 ± 1.56 min; Fig 1f). Both the lysosome and mitochondria are key sites for sensing metabolic stress, and distinct mechanisms for AMPK activity have been identified at these compartments.^20–23^ This is also consistent with previous findings that AMPK activity in response to energy stress is most rapidly induced at the mitochondria and lysosome.^14^ These results suggest that AMPK is rapidly activated at sites of high metabolic flux or local oxidative stress accumulation.

To assess the relationship between oxidative stress and AMPK activity, we quantified cellular oxidation across a range of H₂O₂ concentrations using HyPer7, a genetically encoded hydrogen peroxide biosensor.^24^ We performed this analysis across multiple subcellular compartments, including the cytoplasm, nucleus, mitochondria, and lysosome (Supplementary Fig. 1). We found that in the cytoplasm and nucleus, Hyper7 response generally scaled with increasing H₂O₂ concentration, and AMPK activity exhibited a corresponding dose-dependent response. At the lysosome, the dose-dependent response seemed to reach a maxima, with both ExRai AMPKAR and HyPer7 performing similarly at the medium and high doses of H_2_O_2_ tested, suggesting high sensitivity to oxidative stress at this location. At the mitochondria, we did not observe this dose-dependent response. This could be because the mitochondria is a site of reactive oxygen species formation and is inherently sensitive to oxidative stress. These data support a model in which AMPK activation is regulated by oxidative stress and tuned by the local subcellular microenvironment.

### LKB1 is required for compartmentalized AMPK activation in response to oxidative stress

Maximal AMPK activation under cellular stress is driven by phosphorylation of AMPKɑ1/2 Thr172 by liver kinase B1 (LKB1) and Ca^2+^/calmodulin-dependent protein kinase kinase 2 (CaMKK2).^25^ We previously found that LKB1 has distinct impacts on compartmentalized AMPK activity in response to stimuli including 2-deoxyglucose (2-DG) and the allosteric activator MK-8722.^14^ To determine if upstream kinases have a similar impact on the spatial regulation of AMPK during oxidative stress, we treated LKB1 knockout (KO) MEFs with H₂O₂ and measured ExRai AMPKAR response in the cytoplasm, nucleus, at the mitochondria, and lysosome (Supplemental Fig 2a). In all compartments, LKB1 KO MEFs exhibited suppressed AMPK activity in response to H₂O₂ stimulation, which we confirmed by western blotting for phosphorylation of the downstream AMPK target, acetyl-CoA carboxylase (ACC; e.g. cytoplasm WT ΔR/R_0_=3.21 ± 0.22, LKB1 KO ΔR/R_0_=0.29 ± 0.04, \*\*\*\**p*<0.0001; Fig. 2a–d, Supplemental Fig 2b).^26^ This is in contrast to previous findings that knocking out LKB1 led to location-specific changes in 2-DG or MK-8722 induced AMPK activity.^14^

**Figure 2.**
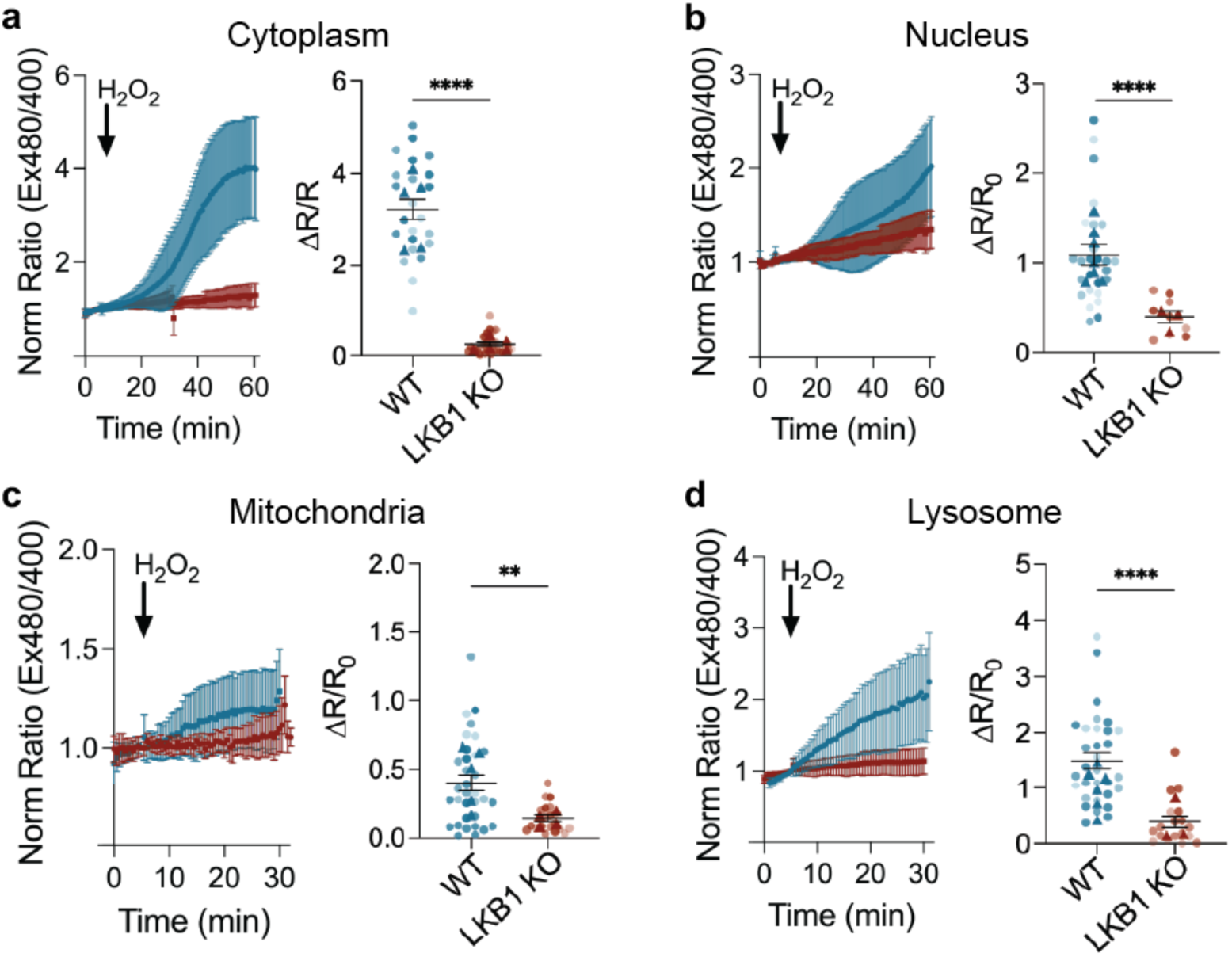
Oxidative stress-induced AMPK activity is dependent on LKB1. **a** Average response of ExRai AMPKAR in WT (blue, reproduced from 1b) and LKB1 KO (dark red, n=25 from five experiments) MEFs treated with H_2_O_2_ (150 µM), along with maximum ratio change (\*\*\*\**p*<0.0001, unpaired t-test, two-tailed). **b** Average response of NLS-ExRai AMPKAR in WT (blue, reproduced from 1c) and LKB1 KO (dark red, n=9 from three experiments) MEFs treated with H_2_O_2_ (150 µM), along with maximum ratio change (\*\*\*\**p*<0.0001, unpaired t-test, two-tailed). **c** Average response of mito-ExRai AMPKAR in WT (blue, reproduced from 1d) and LKB1 KO (dark red, n=21 from four experiments) MEFs treated with H_2_O_2_ (150 µM), along with maximum ratio change (\*\**p*=0.0013, unpaired t-test, two-tailed). **d** Average response of lyso-ExRai AMPKAR in WT (blue, reproduced from 1e) and LKB1 KO (dark red, n=18 from three experiments) MEFs treated with H_2_O_2_ (150 µM, along with maximum ratio change (\*\*\*\**p*<0.0001, unpaired t-test, two-tailed). For all figures, time courses show the mean ± SD, dot plots show the mean ± SEM.

We also examined the role of CaMKK2 in H_2_O_2_-induced AMPK activity using CaMKK2 knockout MEFs^14^ (Supplementary Fig. 2a-b). In the absence of CaMKK2, we observed minimal change in AMPK activity in the cytoplasm (WT ΔR/R_0_=3.21 ± 0.22, CaMKK2 KO ΔR/R_0_=2.40 ± 0.21; *p*=0.4127; Supplemental Fig 2c), nucleus, (WT ΔR/R_0_=1.02 ± 0.11, CaMKK2 KO ΔR/R_0_=1.58 ± 0.23; *p*=0.1167; Supplemental Fig 2d), or lysosome (WT ΔR/R_0_=1.15 ± 0.11, CaMKK2 KO ΔR/R_0_=1.78 ± 0.25; *p*=0.0357 Supplemental Fig 2f), following H₂O₂ treatment. However, mitochondrial AMPK activity was significantly diminished (WT ΔR/R_0_=0.39 ± 0.06, CaMKK2 KO ΔR/R_0_=0.28 ± 0.08, *p*=0.0020; Supplemental Fig 2e), suggesting that CaMKK2 contributes to mitochondrial AMPK activation under oxidative stress (Supplementary Fig. 2g). These results are consistent with previous findings that loss of CaMKK2 diminishes AMPK activity at the mitochondria.^14^ Altogether, these results indicate that LKB1 is required for global AMPK activation in response to H_2_O_2_, while CaMKK2 plays a more localized, compartment-specific role at the mitochondria.

### Phosphorylation of AMPK by upstream kinases is necessary for oxidative stress-induced AMPK activity

Based on our findings that LKB1 is important for AMPK activation throughout the cell under oxidative stress, we hypothesized that AMPK activity in response to oxidative stress is due to phosphorylation of Thr172 on the AMPKɑ subunit. We imaged ExRai AMPKAR in wildtype or AMPKα1/2 KO U2OS cells treated with hydrogen peroxide, finding that AMPKɑ1/2 KO U2OS cells had minimal AMPK activity compared to WT (WT ΔR/R_0_=2.58 ± 0.15, α1/2 KO ΔR/R_0_=0.28 ± 0.07, ****p<0.0001; Fig. 3a, Supplemental Fig 3a). We then rescued AMPK activity by co-expressing ExRai AMPKAR alongside mScarlet-AMPKα2 or mScarlet-AMPKα2 T172A, which cannot be phosphorylated by upstream kinases, and measured AMPK activity using ExRai AMPKAR (Fig. 3b). Rescuing AMPK expression using mScarlet-AMPKα2 expressed in AMPKα1/2 KO U2OS cells partially restored AMPK activity (ΔR/R_0_=1.08 ± 0.13) as compared to AMPKα1/2 KO U2OS (*p*<0.0001). Conversely, mScarlet-AMPKα2 T172A could not restore AMPK activity (ΔR/R_0_=0.21 ± 0.01, *p*=0.3815). This demonstrates that upstream phosphorylation is required for oxidative stress-induced AMPK activity.

**Figure 3:**
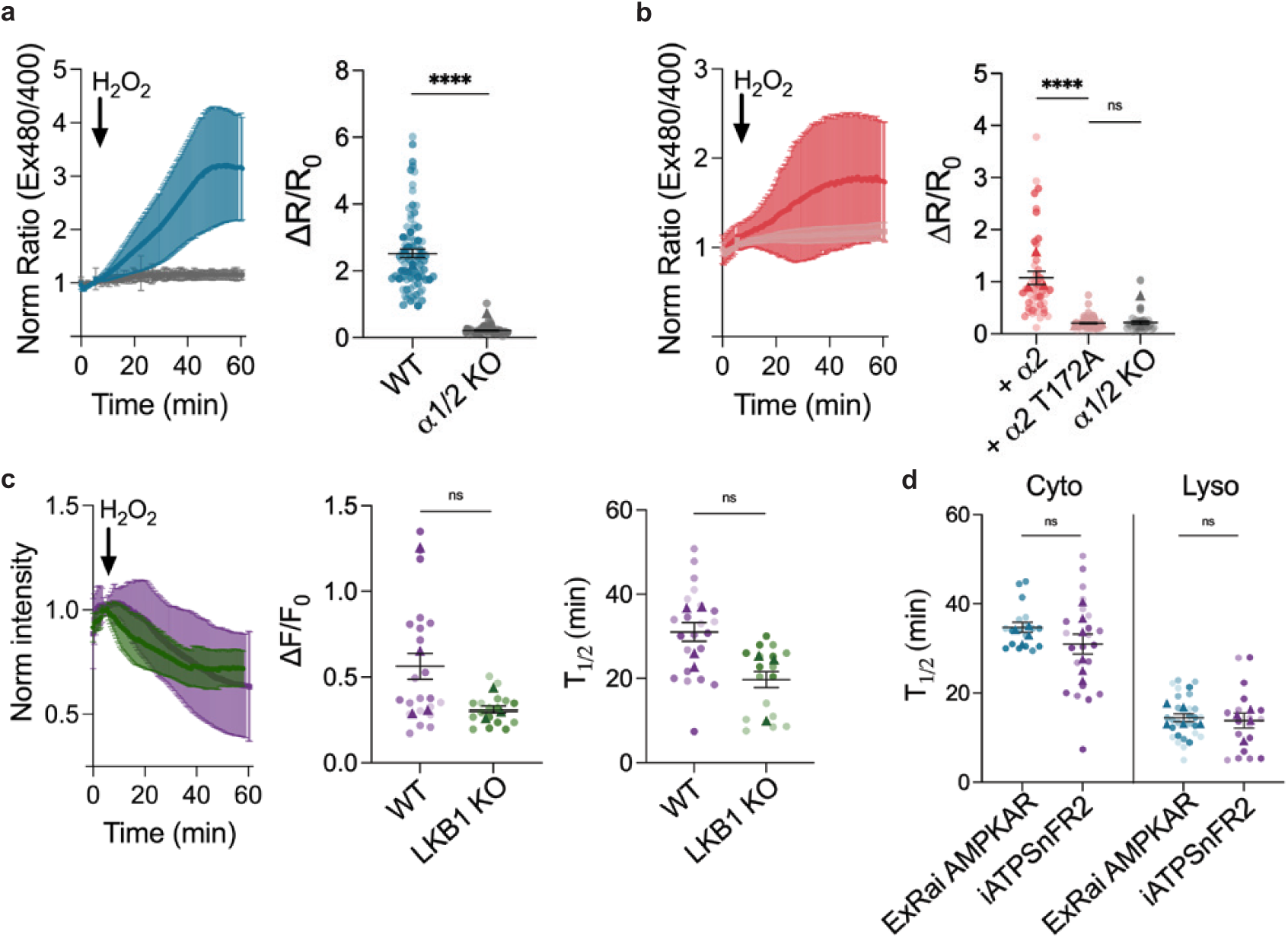
Oxidative stress-induced AMPK activity requires phosphorylation by an upstream kinase. **a** Average response of ExRai AMPKAR in WT U2OS (blue, n=81 from three experiments) and AMPKα1/2 KO U2OS (grey, n=59 from three experiments) treated with H_2_O_2_ (150 µM), along with maximum ratio change (****p<0.0001, unpaired t-test, two-tailed). **b** Average response of ExRai AMPKAR in AMPKα1/2 KO U2OS with AMPKα2 (pink, n=43 from three experiments) and with AMPKα2 T712A (light pink, n=89 from three experiments) treated with H_2_O_2_ (150 µM), along with maximum ratio change (****p<0.0001, ns=0.3815, unpaired t-test, two-tailed). AMPKɑ1/2 KO U2OS ΔR/R_0_ replotted from a. **c** Average response of iATPSnFR2 in WT (purple, n=22 cells from four experiments) and LKB1 KO (green, n=17 cells from three experiments) MEFs treated with H_2_O_2_ (150 µM), along with maximum intensity change (ns=0.4000, unpaired t-test, two-tailed) and *t*_1/2_ (ns=0.1667, unpaired t-test, two tailed). **d** *t*_1/2_ of ExRai AMPKAR (blue, n=23 cells from five experiments), and iATPSnFR2 (purple, n=22 cells from four experiments) in WT MEFs treated with H_2_O_2_ (150 µM) (ns=0.2371, unpaired *t*-test, two-tailed), alongside *t*_1/2_ of lyso ExRai AMPKAR (blue, n=23 cells from five experiments), and iATPSnFR2 (purple, n=22 cells from four experiments) in WT MEFs treated with H_2_O_2_ (150 µM, ns=0.2371, unpaired *t*-test, two-tailed). For all figures, time courses show the mean ± SD, dot plots show the mean ± SEM

AMPK activation by hydrogen peroxide has been reported to occur due to depletion of ATP.^15,16^ However, our data suggests LKB1 is necessary for AMPK activity in response to oxidative stress. To determine if changes in ATP drive oxidative stress-induced AMPK activity, we imaged ATP dynamics in the cytoplasm of WT and LKB1 KO MEFs treated with hydrogen peroxide with the ATP biosensor iATPSnFR2^27^ (Fig. 3c). While we observed differences in AMPK activity between WT and LKB1 KO MEFs (Fig. 2), we found WT and LKB1 KO MEFs had similar changes in ATP dynamics, both with the level of ATP depletion (WT ΔF/F_0_=0.56 ± 0.08, LKB1 KO ΔF/F_0_= 0.31 ± 0.02, *p*=0.4000) and kinetic response (WT *t*_1/2_= 31.04 ± 2.24 min, LKB1 KO *t*_1/2_= 19.72 ± 1.92 min, *p*=0.1667).

When comparing the t_1/2_ of ExRai AMPKAR to iATPSnFR2 in the cytoplasm of WT MEFs, we found AMPK activation occurs with similar kinetics as ATP depletion (ExRai AMPKAR t_1/2_=36.52 ± 1.56, iATPSnFR2 t_1/2_=31.04 ± 2.24 min, *p*=0.2371; Fig. 3d). We extended this analysis to the lysosome, a compartment increasingly recognized as a privileged site for integrating metabolic and redox signaling. We similarly observed no significant difference in kinetics between local AMPK activation and ATP depletion (Lyso-ExRai AMPKAR t_1/2_=14.48 ± 0.86, Lyso-iATPSnFR2 t_1/2_=13.84 ± 1.69 min, *p*=0.5942; Fig. 3d). We performed cross-correlation analysis and found that ExRai AMPKAR response and iATPSnFR2 responses were correlated, indicated by peak lag at time 0 (Supplemental Fig 3b). Together, our data suggest a model where oxidative stress-induced AMPK activity is driven by local upstream kinase signaling and local ATP depletion dictates the kinetics of AMPK activity. This supports previous findings that upstream phosphorylation provides the potential for high AMPK activity, but the kinetics of activation are dynamically controlled by the AMP/ATP ratio, which rapidly dictates the conformational state of the kinase domain by modulating the interaction of the ɑ subunit autoinhibitory domain.^28^ Importantly, we find that the local environment (e.g. localized ATP depletion) seems to drive the kinetics of compartmentalized AMPK activity.

### Spatial AMPK activity is minimally induced by oxidative stress in LKB1-null cancer cell lines

LKB1 is a tumor suppressor that is mutated or expression is lost in approximately 10-30% of non-small cell lung cancers (NSCLCs) and cervical cancers.^29^ For NSCLC tumors harboring KRAS mutations, LKB1 loss defines a highly aggressive subtype marked by increased metabolic stress and poor therapeutic response.^30,31^ Similarly, in cervical cancers, loss of LKB1 promotes increased tumor cell metabolism and increases reactive oxygen species.^32^ Given the role of LKB1 in activating AMPK under oxidative stress, we hypothesized that LKB1-null cancers would have suppressed spatial AMPK activity in response to H_2_O_2_. We used ExRai AMPKAR to measure AMPK activity in various LKB1-null cancer cell lines, including HeLa cells, and KRAS-mutant NSCLC cell lines (A549, H460, and H1355). Stimulation with H_2_O_2_ resulted in suppressed AMPK activation in all LKB1-deficient cell lines, while co-expression of LKB1-mCherry with ExRai AMPKAR enhanced AMPK activity in each cell line (Fig. 4a,d,g,j). To assess spatial AMPK activity in these cells, we used lysosome-targeted ExRai AMPKAR and measured AMPK response to H_2_O_2_. We observed some lysosomal AMPK activation in these cells, which was significantly enhanced with LKB1-mCherry rescue in HeLa, A549, H1355, and H460 cells (Fig. 4b,e,h,k). These results confirm that LKB1 is needed for maximal AMPK activation in response to oxidative stress in these cancer cell lines.

**Figure 4.**
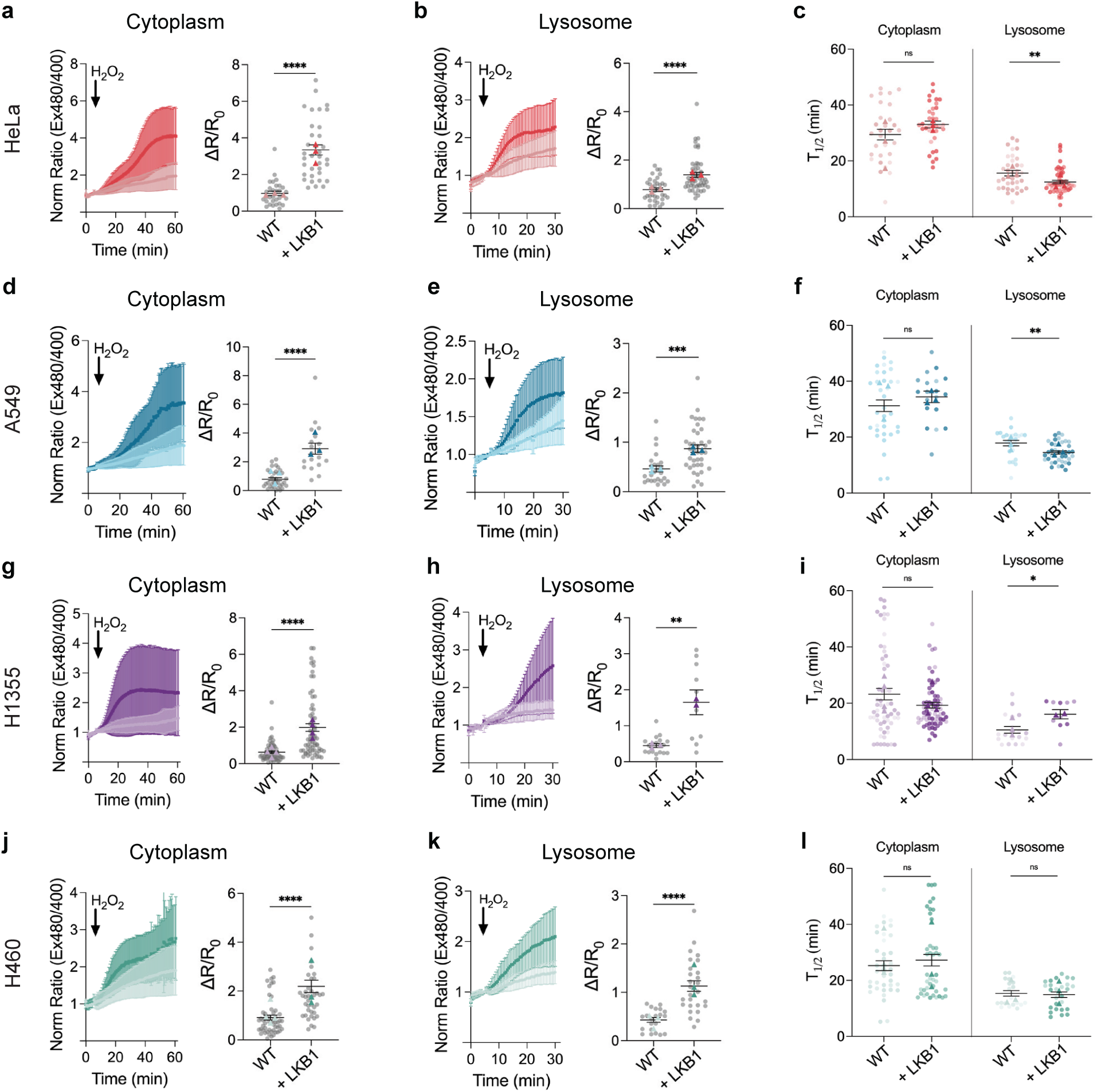
LKB1 drives spatial AMPK activity in response to oxidative stress in cancer cells. **a** Average response in HeLa cells expressing ExRai AMPKAR (light pink, n=31 cells from three experiments) or co-expressing with mCherry-LKB1 (dark pink, n=35 cells from three experiments) treated with H_2_O_2_ (150 µM), along with maximum ratio change (\*\*\*\**p*<0.0001, unpaired *t*-test, two-tailed). **b** Average response in HeLa cells expressing lyso-ExRai AMPKAR in HeLa cells (light pink, n=35 cells from three experiments) or co-expressing with mCherry-LKB1 (dark pink, n=52 cells from three experiments) treated with treated with H_2_O_2_ (150 µM), along with maximum ratio change (\*\*\*\**p*<0.0001, unpaired *t*-test, two-tailed). **c** *t*_1/2_ in HeLa cells expressing cytoplasmic ExRai AMPKAR (light pink), or co-expressing with mCherry-LKB1 (dark pink), lyso-ExRai AMPKAR (light pink) or co-expressing with mCherry-LKB1 (dark pink) treated with treated with 150 µM H_2_O_2_ (ns=0.1163, \**p*=0.0069, unpaired *t*-test, two-tailed). t_1/2_ quantified from a and b. **d** Average response in A549 cells expressing ExRai AMPKAR (light blue, n=34 cells from three experiments) or co-expressing with mCherry-LKB1 (dark blue, n=19 cells from three experiments) treated with H_2_O_2_ (150 µM), along with maximum ratio change (\*\*\*\**p*<0.0001, unpaired *t*-test, two-tailed). **e** Average response in A549 cells expressing lyso-ExRai AMPKAR in A549 cells (light blue, n=25 cells from three experiments) or co-expressing with mCherry-LKB1 (dark blue, n=38 cells from three experiments) treated with treated with H_2_O_2_ (150 µM), along with maximum ratio change (\*\*\**p*=0.0001, unpaired *t*-test, two-tailed). **f** *t*_1/2_ in A549 cells expressing cytoplasmic ExRai AMPKAR in A549 cells (light blue), co-expressing with LKB1 (dark blue), lyso-ExRai AMPKAR (light blue) or co-expressing with mCherry-LKB1 (dark blue) treated with treated with 150 µM H_2_O_2_ (ns=0.4341, \*\**p*=0.0015, unpaired *t*-test, two-tailed). t_1/2_ quantified from d and e. **g** Average response in H1355 cells expressing ExRai AMPKAR (light purple, n=59 cells from four experiments) or co-expressing with mCherry-LKB1 (dark purple, n=66 cells from four experiments) treated with H_2_O_2_ (150 µM), along with maximum ratio change (\*\*\*\**p*<0.0001, unpaired *t*-test, two-tailed). **h** Average response in H1355 cells expressing lyso-ExRai AMPKAR (light purple, n=19 cells from three experiments) or co-expressing with mCherry-LKB1 (dark purple, n=15 cells from three experiments) treated with treated with H_2_O_2_ (150 µM), along with maximum ratio change (\*\**p*=0.0014, unpaired *t*-test, two-tailed). **i** *t*_1/2_ in H1355 cells expressing cytoplasmic ExRai AMPKAR in H1355 (light purple) or co-expressing with mCherry-LKB1 (dark purple), lyso-ExRai AMPKAR (light purple) or co-expressing with mCherry-LKB1 (dark purple) treated with 150 µM H_2_O_2_ (ns=0.0655, \**p*=0.0162, unpaired *t*-test, two-tailed). t_1/2_ quantified from g and h. **j** Average response in H460 cells expressing ExRai AMPKAR (light teal, n=53 cells from four experiments) or co-expressing with mCherry-LKB1 (dark teal, n=42 cells from three experiments) treated with H_2_O_2_ (150 µM), along with maximum ratio change (\*\*\*\**p*<0.0001, unpaired *t*-test, two-tailed). **k** Average response in H460 cells expressing lyso-ExRai AMPKAR in H460 (light teal, n=21 cells from three experiments) or co-expressing with mCherry-LKB1 (dark teal, n=26 cells from three experiments) treated with treated with H_2_O_2_ (150 µM), along with maximum ratio change (\*\*\*\**p*<0.0001, unpaired *t*-test, two-tailed). **l** *t*_1/2_ in H460 cells expressing ExRai AMPKAR (light teal) or co-expressing with mCherry-LKB1 (dark teal), lyso-ExRai AMPKAR (light blue) or co-expressing with mCherry-LKB1 (dark teal) treated with treated with 150 µM H_2_O_2_ (ns>0.6183, unpaired *t*-test, two-tailed). t_1/2_ quantified from j and k. For all figures, time courses show the mean ± SD, dot plots show the mean ± SEM.

Previously, we found that loss of LKB1 resulted in slower accumulation of AMPK activity in response to 2-DG or MK-8722, as measured by ExRai AMPKAR *t_1/2_*.^14^ As these LKB1-null cells had some measurable lysosomal AMPK activity in response to H_2_O_2_, we hypothesized that LKB1 rescue would increase accumulation of lysosomal AMPK activity. We quantified the kinetics of lysosomal ExRai AMPKAR response in these cell lines and found that in HeLa and A549 cells, lysosomal AMPK activation occurred more rapidly than in the cytoplasm when LKB1 expression was restored (Fig. 4c,f,I,l). However, in H460 cells, while re-expression of LKB1 did increase the response of ExRai AMPKAR (ΔR/R_0_=1.14 ± 0.11), the rate that AMPK activity accumulated at the lysosome was not significantly impacted (t_1/2_ = 14.90 ± 0.96 min).

Additionally, in H1355 cells, AMPK activity at the lysosome occurred slower after reexpression of LKB1 (t_1/2_=16.12 ± 1.64 min). For both of these cell lines, this minimal change or increase in t_1/2_ could be reflective of the robust increase in AMPK activity at the lysosome. In the cytoplasm of all cell lines, the t_1/2_ of ExRai AMPKAR was not significantly increased upon LKB1 re-expression, only the magnitude of the response (ΔR/R_0_), consistent with our previous work.^14^ Our results extend this observation to oxidative stress, demonstrating LKB1 regulation of AMPK has location-specific dynamics.

### AMPK regulates cellular response to oxidative stress

AMPK is critical for maintaining cellular homeostasis, with over 100 identified downstream targets throughout the cell.^33^ While previous work has identified the downstream effectors of AMPK in response to energy stress^34,35^ or rising intracellular calcium levels,^36–38^ the downstream signaling targets of AMPK under oxidative stress remain poorly characterized. To identify specific downstream targets of AMPK under oxidative stress, we performed phosphoproteomics using MEFs treated with H_2_O_2_, with or without pre-treatment with the pharmacological inhibitor of AMPK, SBI-0206965 (Supplemental Fig 4a-c). SBI-0206965 has been shown to inhibit AMPK activity,^39^ which we verified by imaging ExRai AMPKAR in MEFs pre-treated with SBI-0206965 before exposure to H_2_O_2_ (Supplemental Fig. 4d).

We identified 251 differentially regulated phosphosites across 174 unique peptides (Fig. 5a). From our significantly altered phosphopeptides, we identified previously known AMPK downstream targets. Of note, we identified AMPK itself (*PRKAA1*, *PRKAB1*), along with signal transducer and activator of transcription 3 (STAT3), acetyl-CoA carboxylase (ACC, *ACACA*), and nuclear factor-erythroid 2-related factor 2 (NRF2, *NFE2L2*), the phosphorylation of which were all downregulated in SBI-0206965 treated cells (Fig. 5b).^40–42^ These downstream effectors have specific subcellular locations, including the cytoplasm, mitochondrial outer membrane, endosomal membranes, and nucleus, highlighting AMPK activity throughout the cell (Supplementary Fig 4e). We estimated kinase activities from the global phosphoproteomics data using prior knowledge networks of kinase-substrate interactions, which indicated AMPK signaling was strongly downregulated in SBI-0206965-treated conditions (Supplemental Fig. 4f).

**Figure 5.**
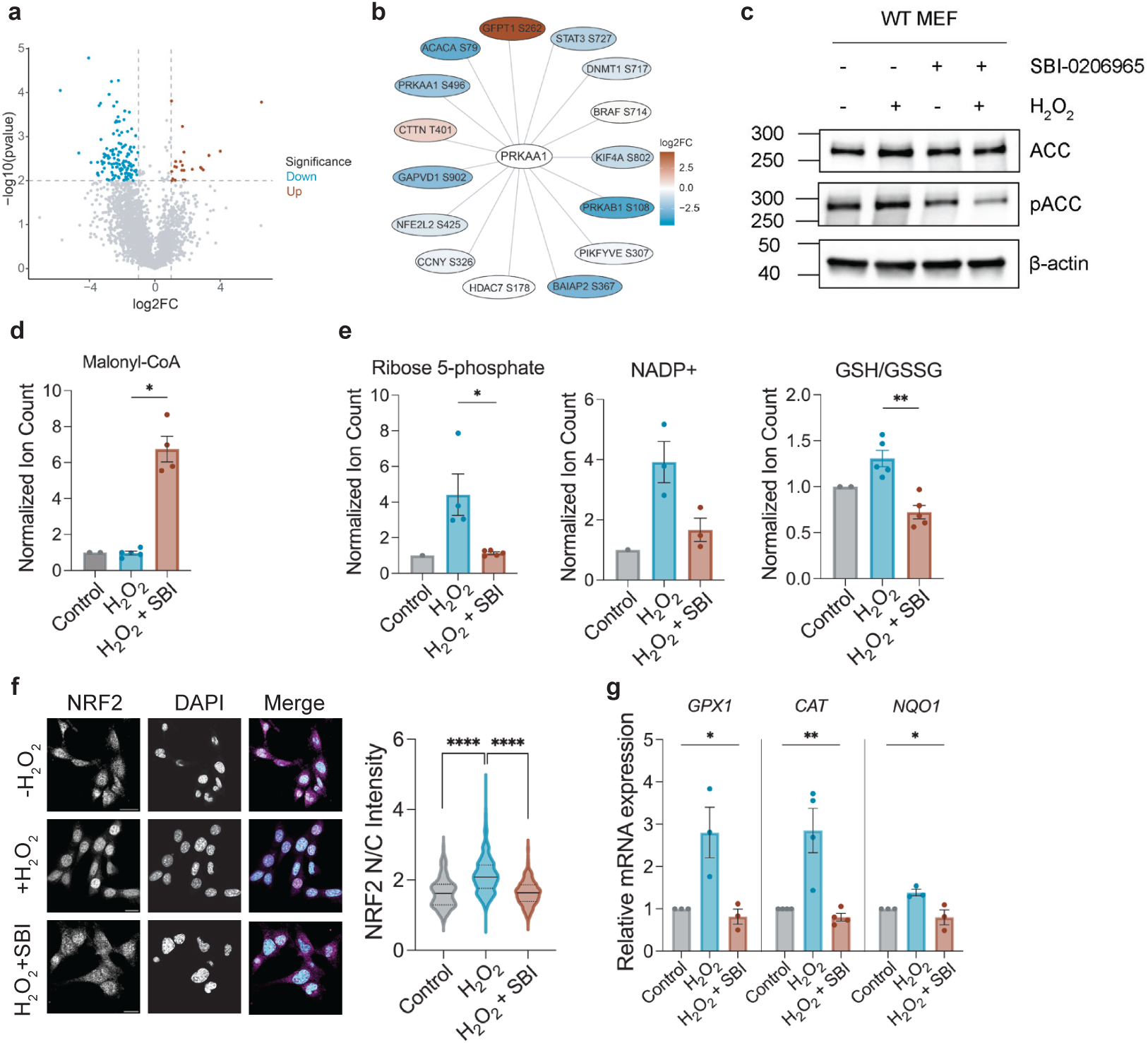
AMPK regulates cellular response to oxidative stress. **a** Volcano plot highlighting changes in phosphopeptides between WT MEFs treated with H_2_O_2_ (150 µM), with and without SBI-0206965 (10 µM). Colored dots depict downregulated (blue, n= 210 peptides) and upregulated (red, n=41 peptides) phosphosites in SBI-0206965 treated conditions at |log2FC > 1| and *p*<0.01 cutoffs. **b** Phosphorylation substrate network for human PRKAA1 (AMPK), comparing WT MEFs treated with H_2_O_2_ (150 µM), with and without SBI-0206965 (10 µM). Proteins are identified using mouse gene names, with corresponding phosphosites identified indicated. Red represents upregulated and blue represents downregulated in SBI-0206965 treated conditions. **c** Representative western blot of WT MEFs treated with H_2_O_2_ (150 µM) with and without SBI-0206965 (10 µM) for total and phospho-acetyl-CoA carboxylase (ACC). Full blots are shown in Source Data **d** Ion count for malonyl-CoA in WT MEFs after treatment with 150 µM H_2_O_2_ (blue) and with pre-treatment with 10 µM SBI-0206965 (red), normalized to vehicle control (grey; n=2 independent experiments with 2-3 technical replicates; \**p*=0.0159, unpaired *t*-test two-tailed). **e** Ion count for ribose 5-phosphate, NADP+, and ratio of GSH to GSSG in WT MEFs treated with 150 µM H_2_O_2_ (blue), and with pre-treatment with10 µM SBI-0206965 (red), normalized to vehicle control (grey; n=2 independent experiments with 2-3 technical replicates; \**p*=0.0159, \*\**p=*0.0079, unpaired *t*-test, two-tailed). **f** Representative immunostaining images of NRF2 (left) and DAPI nuclear stain (center) in WT MEFs treated with H_2_O_2_ (150 µM), with and without pre-treatment with SBI-0206965 (10 µM). Merged images are shown at right. Scale bars represent 20 µm. Violin plot depicts nuclear/cytoplasmic (N/C) NRF2 intensity ratio in WT MEFs treated with 150 µM H_2_O_2_ (blue), vehicle control (grey), and with pre-treatment with SBI-0206965 (red; for all conditions n>195 cells from three independent experiments; \*\*\*\**p*<0.0001, Kruskal-Wallis test). **g** Relative mRNA expression of *GPX1* (GID 14775), *CAT* (NM_009804.2), *NQO1 (*NM_008706) in WT MEFs treated with H_2_O_2_ in WT MEFs after treatment with H_2_O_2_ (150 µM) (blue, n=3 independent experiments with 3 technical replicates), and with pre-treatment with 10 µM SBI-0206965 (red, n=3 independent experiments with three technical replicates), normalized to GAPDH (NM_008084; *\*p*<0.0357, \*\**p*=0.0026) Kruskal-Wallis test).

To validate of our data set, we assessed the phosphorylation of STAT3 S727, which was downregulated in WT MEFs pre-treated with SBI-0206965 prior to H_2_O_2_ treatment. AMPK has been reported to phosphorylate STAT3 at this site.^43^ We found using western blotting that when treated with H_2_O_2_ WT MEFs showed a notable increase in phosphorylated STAT3, where WT MEFs treated with SBI-0206965 showed minimal phosphorylated STAT3 (Supplementary Fig. 4g).

We then assessed the impact of oxidative stress-induced AMPK activity on cell function through functional validation of our top hits. We identified the canonical downstream target of AMPK, ACC, as being regulated by AMPK under oxidative stress. AMPK phosphorylates ACC at S79, and we confirmed ACC S79 phosphorylation was decreased in SBI-0206965 pretreated conditions using western blotting (Fig. 5c). ACC functions to produce malonyl-CoA, the building block of fatty acids, and AMPK phosphorylation of ACC decreases ACC activity and malonyl-CoA levels. To determine if malonyl-CoA levels are decreased when AMPK is activated by H_2_O_2_, we performed metabolomics using MEFs treated with H_2_O_2_, with or without pre-treatment with SBI-0206965 (Fig. 5d, Supplemental Fig. 4h). We found that malonyl-CoA levels were higher in SBI-0206965-pretreated MEFs when compared to non-pretreated MEFs (*p*= 0.0159), suggesting that AMPK downregulates fatty acid biosynthesis during oxidative stress, and when AMPK is inhibited during oxidative stress, ACC is still capable of producing malonyl-CoA.

We also found that ribose 5-phosphate (R5P) levels were higher in SBI-0206965-pretreated MEFs when compared to non-pretreated MEFs (*p*= 0.0159; Fig. 5e, Supplemental Fig 4h). R5P is a key metabolite in the pentose phosphate pathway that supports nucleotide biosynthesis and cellular redox balance through the generation of NADPH. NADPH is oxidized, producing NADP+, to fuel antioxidant systems and maintain the reduced glutathione pool for removal of hydrogen peroxide (Fig 6d).^44,45^ NADPH is also consumed to provide reducing equivalents for fatty acid biosynthesis.^46^ Thus, we examined levels of NADP+. We found that NADP+ levels were significantly higher in MEFs without SBI-0206965 pre-treatment, suggesting that AMPK activation promotes NADPH-utilizing pathways (Fig. 5e). As NADPH is critical for maintaining the glutathione pool, we next assessed the ratio of glutathione (GSH) and oxidized glutathione (GSSG) in WT MEFs treated with hydrogen peroxide, with and without SBI-0206965. The ratio of reduced GSH to GSSG is an indicator of cellular health, with a high ratio indicating robustness in dealing with oxidative stress and a low ratio signaling oxidative damage.^47^ MEFs pretreated with SBI-0206965 prior to H_2_O_2_ introduction had a lower GSH/GSSG ratio compared to non-inhibited cells (*p*=0.0079), suggesting that AMPK is important in maintaining cellular redox homeostasis during oxidative stress (Fig 5e, Supplemental Figure 4i). Together, these data support a model in which AMPK suppresses fatty acid biosysnthesis, which consumes large amounts of NADPH, and promotes the pentose phosphate pathway, observed as R5P accumulation and increased GSH/GSSG ratio, as part of a broader metabolic reprogramming to maintain redox homeostasis during oxidative stress.

**Figure 6.**
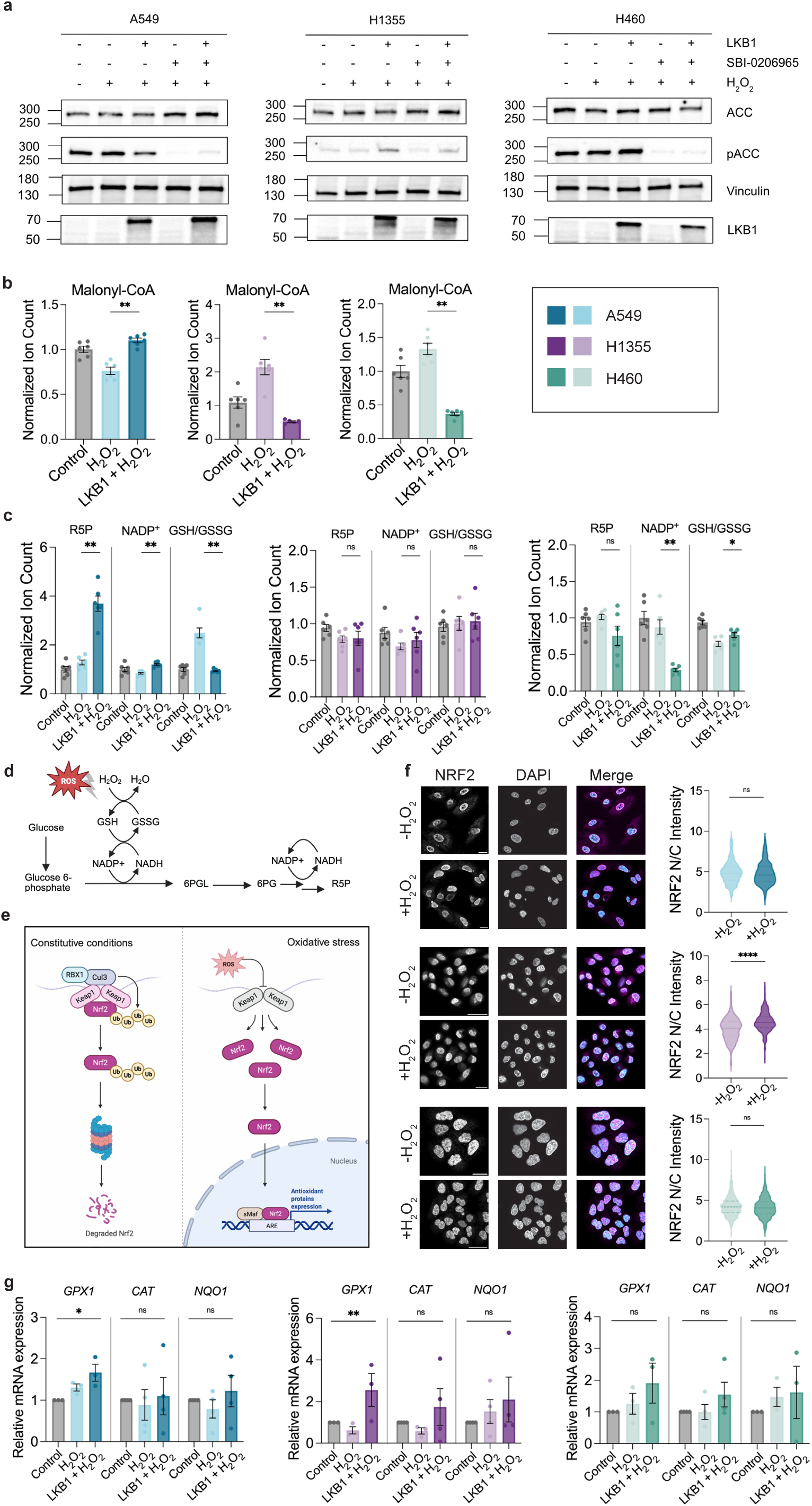
LKB1-AMPK signaling axis is disrupted in KEAP1-mutant cancer cells. **a** (From left to right) Representative western blots of A549, H1355, and H460 cells treated with H_2_O_2_ (150 µM), with and without SBI-0206965 (10 µM) and/or LKB1 re-expression for total and phospho-acetyl-CoA carboxylase (ACC). Full blots are shown in Source Data. **b** Ion count for malonyl-CoA in A549 (blue), H1355 (purple), and H460 (teal) cells treated with H_2_O_2_ (150 µM), with and without LKB1 re-expression, normalized to vehicle control (grey, n=1 independent experiment with 6 technical replicates; \**p*<0.0195, \*\**p*<0.0043, unpaired *t*-test, two-tailed). **c** Ion count for ribose 5-phosphate (R5P), NADP+, and ratio of GSH to GSSG in A549 (blue), H1355 (purple), and H460 (teal) cells treated with H_2_O_2_ (150 µM), with and without LKB1 re-expression, normalized to vehicle control (grey, n=1 independent experiment with 6 technical replicates; ns≥0.2523 \**p*<0.0476, \*\**p*<0.0022, unpaired *t*-test, two-tailed). **d** Schematic model of pentose phosphate pathway- dependent NADH and glutathione regeneration during oxidative stress. **e** Schematic model of KEAP1-mediated NRF2 regulation under basal and oxidative stress conditions.’ **f** (From top to bottom) Representative immunostaining images of NRF2 (left) and DAPI nuclear stain (center) in A549, H1355, and H460 cells treated with H_2_O_2_ (150 µM). Merged images are shown at right. Scale bars represent 20 µm. Violin plot depicts nuclear/cytoplasmic (N/C) NRF2 intensity ratio in HeLa (pink), A549 (blue), H1355 (purple), and H460 (teal) cells (for all conditions n≥116 cells from three independent experiments; ns≥0.1508, \*\*\*\**p*<0.0001, unpaired t-test, two-tailed). **g** Relative mRNA expression of *GPX1* (GID 14775), *CAT* (NM_009804.2), and NQO1 (NM_008706) in HeLa (pink), A549 (blue), H1355 (purple), and H460 (teal) cells treated with H2O2 (150 µM), with and without LKB1 re-expression, normalized to GAPDH (n=3 independent experiments with 3 technical replicates; ns≥0.1333, \**p*=0.0250, \*\**p*=0.0036, Kruskal-Wallis test).

We finally examined NRF2, a transcription factor central to the antioxidant response. AMPK is known to modulate NRF2 signaling through phosphorylation at Ser558, which blocks NRF2 removal from the nucleus and promotes transcription of various genes involved in the antioxidant stress response, including glutathione peroxidase (*GPX1*), catalase (*CAT*), and NAD(P)H quinone dehydrogenase 1 (*NQO1*).^48,49^ As we identified suppressed NRF2 S558 phosphorylation when AMPK activity was inhibited during oxidative stress, we hypothesized that AMPK-NRF2 axis would regulate the expression of these genes in response to oxidative stress (Fig 5b). To test this, we first examined NRF2 localization by immunostaining WT MEFs treated with H_2_O_2,_ with or without pre-treatment with SBI-0206965, and found that AMPK activity was required for efficient NRF2 nuclear translocation (Fig. 5f).

To further assess the functional consequences of this regulation, we performed RT-qPCR on NRF2 target genes *GPX1*, *CAT*, and *NQO1* in WT MEFs treated with H_2_O_2_, with or without SBI-0206965 pretreatment. mRNA expression of these NRF2 target genes was elevated in non-inhibited cells, while in inhibited cells the expression of each gene was similar to control (Fig 5g). Together, these results confirm that under oxidative stress, AMPK phopshorylates NRF2, promoting the nuclear relocalization of NRF2 and enhanced transcriptional activity. More broadly, we demonstrate that under oxidative stress, AMPK regulates metabolism and expression of genes involved in the antioxidant response to maintain redox homeostasis.

### KEAP1 mutation results in incomplete rescue of LKB1-AMPK signaling axis in response to oxidative stress in KLK-mutant NSCLCs

We finally sought to determine how our identified mechanism for how AMPK rewires cell function under oxidative stress is perturbed in KRAS/LKB1/KEAP1 (KLK)-mutant NSCLC cell lines (A549, H460, and H1355), which are reported to exhibit high resistance to chemotherapeutics that induce oxidative stress.^50^ We first assessed ACC phosphorylation as a readout of AMPK activity. Consistent with our previous imaging data (Fig 4), we found these cells exhibited minimal ACC phosphorylation (pACC) in response to H_2_O_2_, while LKB1 reexpression restored pACC levels in H1355 and H460 cells, with minmal rescue in A549 cells (Fig. 6a).

To determine how KLK mutations affects metabolic pathways downstream of AMPK, we performed metabolomics on these cells in response to H_2_O_2_, with and without re-expression of LKB1. In H1355 and H460 cells, malonyl-CoA levels decreased under oxidative stress in an LKB1-dependent manner, consistent with AMPK-mediated inhibition of fatty acid synthesis (Fig. 6b). However, A549 cells displayed a distinct response, with decreased malonyl-CoA levels under LKB1 deficient conditions and elevated malonyl-CoA levels under LKB1 expression, cosnsistent with ACC phosphorylation in these cells (Fig. 6a, b). A549 cells are reported to have high resistance to oxidative stress, and our data suggest that this may be due to compensatory metabolic rewiring independent of LKB1-AMPK regulation.^51,52^

We next examined metabolites within the pentose phosphate pathway (PPP) to assess redox-supporting metabolism. We found that NSCLC cell lines exhibited heterogeneous responses, with variable changes in PPP intermediates and glutathione metabolism across conditions (Fig. 6c,d). A549 cells showed a partial restoration of redox metabolites following LKB1 re-expression, while H460 and H1355 cells displayed minimal inconsistent responses, suggesting disruption of coordinated metabolic regulation downstream of AMPK.

To further assess antioxidant responses, we examined NRF2 activity in these cell lines. Under basal conditions, NRF2 is sequestered in the cytoplasm by KEAP1 and targeted for ubiquitin-mediated degredation. During oxidative stress, this interaction is disrupted, allowing NRF2 to accumulate and translocate to the nucleus where it induces antioxidant gene expression (Fig. 6e).^53^ Notably, the NSCLC lines used here harbor KEAP1 mutations, which we hypothesized would impair NRF2 degredation and result in constituitive NRF2 stabilization and nuclear localization. To test this, we performed immunostaining for NRF2 localization following H_2_O_2_ treatment. In KLK-mutant NSCLCs, NRF2 exhibited high baseline nuclear localization that was largely insensitive to oxidative stress, consistent with constiutive NRF2 activation downstream of KEAP1 mutation (Fig. 6f). Only H1355 cells had a small but significant increase in nuclear NRF2 levels. We next performed RT-qPCR on NRF2 target genes *GPX1, CAT,* and *NQO1*, finding that these cell lines displayed minimal or inconsistent transcriptional resposnes to oxidative stress or LKB1 restoration, further supporting our hypothesis that constituitive NRF2 activation in KLK-mutant cells limits dynamic redox responsiveness (Fig. 6g).

## Discussion

In the present work, we use an AMPK biosensor to reveal that AMPK exhibits distinct spatiotemporal activation patterns in response to hydrogen peroxide (H_2_O_2_). This activation is dependent on the upstream kinase LKB1 and context-dependently CaMKK2, suggesting a model in which upstream kinases locally control AMPK responses to oxidative stress.

Furthermore, we find that LKB1-deficient cancer cells demonstrate suppressed AMPK activity in response to hydrogen peroxide, potentially contributing to their high reported levels of oxidative stress.^54,55^ These results support previous reports implicating LKB1 in the regulation of AMPK activity in response to oxidative stress, and highlight new dimensions of compartmentalized AMPK regulation.^9,19^ By extending these findings to lung and cervical cancer cell lines, we demonstrate how this spatial regulation of AMPK is maintained in healthy cells and perturbed in disease.

Using organelle-targeted ExRai AMPKAR, we observed robust AMPK activation in response to H_2_O_2_ at all subcellular locations tested, with the most rapid responses occurring at the lysosome and mitochondria. These data extend recent findings that AMPK activation is spatially encoded^14,56^ and point to a model in which the lysosome and mitochondria serve as privileged sites for stress signal integration.^57,58^ Lysosomal AMPK pools have previously been implicated in nutrient sensing and autophagy regulation,^20^ while mitochondria are both a major site of reactive oxygen species (ROS) production and a hub for AMPK-mediated mitochondrial quality control.^21,22^ The spatiotemporal differences we observed suggest that AMPK activity is not uniform throughout the cell but is shaped by the local signaling context and subcellular redox landscape.

Activation of AMPK by oxidative stress has been widely reported in the literature, with multiple mechanisms of activation proposed. Early studies proposed that oxidative stress activates AMPK by depleting ATP, thereby increasing the AMP/ATP ratio.^15,16^ Others have suggested that oxidative stress may activate AMPK directly via cysteine oxidation on the α subunit or upstream kinases.^11,17–19^ Our results demonstrate that oxidative stress-induced AMPK activity is suppressed in LKB1 knockout cells, even though ATP depletion still occurs (Fig. 2; Fig. 3c), and CaMKK2 deletion only impacted AMPK activity at the mitochondria (Supplemental Fig. 2). We further demonstrate that phosphorylation by an upstream kinase is required as AMPKα2 T172A was insufficient in restoring AMPK response in U2OS AMPKα1/2 KO cells treated with hydrogen peroxide (Fig. 3b). While we cannot eliminate the possibility that LKB1 and/or AMPK is directly oxidized by H_2_O_2_ under our conditions, our results suggest LKB1 is the primary regulator of oxidative stress-induced AMPK activity, with local ATP depletion driving the kinetics of AMPK activity during oxidative stress. Taken together with our spatial data, this supports a model in which, under oxidative stress, upstream kinase signaling defines the capacity for local AMPK activation, while the local metabolic and energetic state governs its temporal dynamics across subcellular compartments.

This mechanism has particular relevance in many cancer cells, including NSCLCs, where LKB1 is frequently inactivated or lost. KLK NSCLCs are among the most aggressive molecular subtypes and are characterized by elevated oxidative stress and altered mitochondrial metabolism.^30,31^ In A549, H460, and H1355 NSCLC cells, which are KLK NSCLCs, we observed re-expressing LKB1 enhanced AMPK activation following oxidative stress, suggesting these cancer cell lines have suppressed AMPK activity during oxidative stress (Fig 4). These data suggest that tumor cells lacking LKB1 may be unable to activate protective AMPK signaling under oxidative stress, which may present a therapeutic vulnerability, particularly as we find AMPK is involved in regulating the cellular antioxidant response.

To understand downstream responses to oxidative-stress induced AMPK activity, we used a multi-‘omics approach. We found that AMPK regulates a variety of cell functions in response to oxidative stress, including fatty acid biosynthesis and expression of genes involved in the oxidative stress response. Notably, these targets spanned diverse subcellular compartments, including the cytosol, lysosome, mitochondria, and nucleus, indicating that AMPK is functioning throughout the cell to regulate the oxidative stress response (Supplemental Fig 4e). Of note, inhibition of AMPK increased malonyl-CoA levels via dephosphorylation of ACC, suggesting that under oxidative stress, AMPK regulates fatty acid biosynthesis by phosphorylating ACC. When AMPK is inhibited, we measured a large accumulation of malonyl-CoA, surpassing control. While not investigated here, our results here suggest that fatty acid synthase could be inhibited under oxidative stress independent of AMPK, resulting in accumulation of malonyl-CoA. As fatty acid biosynthesis consumes 14 NADPH for every one molecule of palmitic acid produced, and NADPH is used to reduce GSSH to GSH, inhibition of fatty acid biosynthesis by AMPK could be an adaptive response to redirect cellular reducing equivalents to neutralizing oxidative species. Consistent with this, we found that R5P, NADP+, and the GSH/GSSG ratio were decreased in SBI-0206965 pre-treated MEFs, suggesting that AMPK promotes antioxidant capacity through NADPH availability via the pentose phosphate pathway. Supporting the role of AMPK in regulating the cellular oxidative stress response, we find that AMPK activation increases the expression of genes involved in this response under the regulation of NRF2 (Fig. 5f).

Extending these findings to NSCLCs, we observe that this coordinated metabolic and transcriptional response are differentially regulated in KLK-mutant NSCLC cell lines. Regulation of fatty acid metabolism remained generally consistent with AMPK dependency, as restoration of LKB1 re-established ACC phosphorylation or reduced malonyl-CoA levels in most cell lines examined (Fig. 6a,b). In contrast, antioxidant transcriptional responses were highly heterogeneous across NSCLC cell lines. These cells harbor KEAP1 mutations, which we found to constituitively stabilize NRF2 and maintain its nuclear localization independent of upstream oxidative stress signaling (Fig. 6e). As a result, antioxidant gene expression appears to become uncoupled from AMPK activity, potentially explaining why these tumors tolerate the chronically elevated oxidative stress associated with KRAS-driven metabolism (Fig. 6g).^59^ This constitutive NRF2 activity may allow KLK-mutant NSCLCs to maintain antioxidant defenses even in the absence of coordinated AMPK signaling, thereby contributeing to their well-described resistance to oxidative stress and therapeutic targeting.^60,61^ These results illustrate that AMPK serves not only as a master regulator of energy stress, but also as a key node linking redox sensing to cellular adaptations to environmental stressors.

This work utilizes a novel approach for uncovering compartmentalized AMPK signaling by combing the imaging-based measurement of spatiotemporal dynamics of AMPK activity with a multi-‘omics approach. Using spatially resolved imaging strategies, we uncover a mechanism by which oxidative stress activates AMPK via LKB1 in a location-specific manner with kinetics driven by local ATP depletion, resolving a long-standing debate in the field. Our approach of using biosensors for AMPK and ATP enabled this precise spatial and temporal investigation into mechanisms for dynamic, compartmentalized regulation of AMPK activity. Integrating this with phosphoproteomic and metabolic analysis, we define a downstream network of AMPK-regulated targets specific to redox stress, revealing metabolic programs that prioritize redox balance over biosynthesis. We uniquely combine subcellular single-cell measurements of AMPK activity and metabolite dynamics with ensemble measurements of the phosphoproteome and metabolome for a holistic determination of the mechanism of and role for AMPK activity under oxidative stress. This approach can be used more broadly to study the compartmentalization of other signaling networks and metabolic processes.

Taken together, our findings demonstrate that oxidative stress-induced AMPK activation is spatially encoded and upstream kinase-dependent. We have developed a model where, under oxidative stress, AMPK activity exhibits location-specific activity dynamics, regulated by LKB1 and kinetically influenced by ATP depletion. AMPK then regulates the cellular oxidative stress response. This work corroborates previous findings that subcellular context influences AMPK activity and uncovers downstream consequences of AMPK activity under oxidative stress. The ability to measure AMPK activity in real time and in specific cellular compartments using genetically encoded biosensors provides powerful insight into the logic of stress response signaling. By linking spatially resolved AMPK activity to downstream pathways, our data highlight how stress adaptation is likely rewired in LKB1-deficient cancers. Ultimately, decoding these spatial signaling patterns may yield new strategies for targeting redox and metabolic vulnerabilities in cancer and other diseases.

## Methods

### Materials

Hydrogen peroxide (Fisher Scientific H325-500) was diluted in molecular biology-grade water (Thermo Scientific Cat # BP2819100). SBI-0206965 (Selleck Chemicals Cat # S7885) was diluted in DMSO. For antibodies, the following were used: from BioRad goat anti-rabbit IgG HRP conjugate (1706515) and goat anti-mouse IgG HRP conjugate (1706516) were diluted to 1:10,000; from Cell Signaling Technology, LKB1 (3047S), AMPKɑ (2532S), p-ACC S79 (11818S), ACC (3676S), cofilin (5175S), STAT3 (9139T0), p-STAT3 S727 (9134T) were diluted 1:1000, and β-actin (3700S) was diluted 1:1000; from Santa Cruz Biotech CaMKK2 (sc-100364) was diluted 1:250.

### Plasmids

All primers used for molecular cloning are listed in Supplementary Table 1. mScarlet-AMPKα2 T172A was generated via QuikChange site-directed mutagenesis using Phusion High Fidelity DNA polymerase (Thermo Scientific Cat # F530L) and primers 1-2.^62^ Following PCR amplification, the reaction product was treated with Dpn1 (Thermo Scientific Cat # FD1703) to degrade template DNA and directly transformed into *E. coli* competent cells. Successful clones were confirmed by Sanger sequencing (Genewiz) or whole plasmid sequencing (Plasmidsaurous).

pcDNA3-ExRai-AMPKAR was a gift from Jin Zhang (Addgene plasmid #192446; http://n2t.net/addgene:192446; RRID:Addgene_192446). pcDNA3-Nuc-ExRai-AMPKAR was a gift from Jin Zhang (Addgene plasmid #192450; http://n2t.net/addgene:192450; RRID:Addgene_192450). pcDNA3-Lyso-ExRai-AMPKAR was a gift from Jin Zhang (Addgene plasmid #192449; http://n2t.net/addgene:192449; RRID:Addgene_192449). pcDNA3-Mito-ExRai-AMPKAR was a gift from Jin Zhang (Addgene plasmid #192448; http://n2t.net/addgene:192448; RRID:Addgene_192448). pcDNA3-ExRai-AMPKAR(T/A) was a gift from Jin Zhang (Addgene plasmid #192447; http://n2t.net/addgene:192447; RRID:Addgene_192447). pEGFP-C1-mScarlet-AMPKα2 was a gift from Jin Zhang (Addgene plasmid #192451; http://n2t.net/addgene:192451; RRID:Addgene_192451). pcDNA3-mCherry-LKB1(WT) was a gift from Jin Zhang (Addgene plasmid #84639; http://n2t.net/addgene:84639; RRID:Addgene_84639). pAAV.CAG.iATPSnFR2.A95A.A119L.HaloTag was a gift from Jonathan Marvin (Addgene plasmid #209653; http://n2t.net/addgene:209653; RRID:Addgene_209653). pCS2+MLS-HyPer7 was a gift from Andrew Wojtovich (Addgene plasmid #136470; https://n2t.net/addgene:136470; RRID: Addgene_136470).

### Cell culture and transfection

WT MEFs, LKB1 KO MEFs, and CaMKK2 MEFs were previously described.^14^ HeLa cells were purchased from ATCC (CRM-CCL-2), A549, U2OS, and AMPKα1/2 KO U2OS cells were provided by Reuben Shaw, and H460 and H1355 were provided by David Shackleford. All cell lines were cultured in Dulbecco’s Modified Eagle Medium (DMEM; Gibco Cat #10566016) supplemented with 10% fetal bovine serum (FBS; Gibco Cat # 26140-079), 1% penicillin-streptomycin (Pen/Strep; Gibco Cat #15140-122), and 1 g/L glucose. CaMKK2 knockout mouse embryonic fibroblasts (MEFs) were supplemented with 2µL/mL puromycin (Fisher Scientific Cat #A1113803). Cells were maintained in humidified HeraCell incubators at 37 °C with 5% CO₂. Cells were kept at low passage number (15-25 passages), and mycoplasma contamination was routinely checked using a PCR mycoplasma detection kit (Fisher Scientific, Cat #AAJ66117AMJ).

For transfection, all cell lines were plated on 35-mm glass-bottom dishes (Cellvis, Cat #d35-14-1.5-n) at 50–70% confluency. Cells were transfected 8–24 h after plating using FuGENE HD or 4K (Promega Cat # E2311; E5911) at a 3:1 FuGENE-to-DNA ratio, following the manufacturer’s instructions. ExRai AMPKAR and complementary biosensors were transfected individually or in combination, as indicated. Cells were imaged 24–48 hr post-transfection.

### Epifluorescence imaging and image analysis

For all live-cell imaging experiments, cells were washed and incubated in Hanks’ Balanced Salt Solution (HBSS; Gibco Cat # 14065-056) buffered with 20 mM HEPES (pH 7.4) and supplemented with 1 g/L glucose for at least 30 min at 37°C prior to imaging. Imaging was performed at 37°C. Hydrogen peroxide was diluted to indicated concentration and added at the marked time points, typically after 5 min.

Time-lapse epifluorescence imaging was performed using a Nikon Eclipse Ti2 inverted microscope equipped with equipped with a CHI60 Plan Fluor 40X Oil Immersion Objective Lens (N.A. 1.3, W.D. 0.2 mm, F.O.V. 25mm; Nikon), a Spectra III UV, V, B, C, T, Y, R, nIR light engine featuring 380/20, 475/28, and 575/25 LEDs, Custom Spectra III filter sets (440/510/575 and 390/475/555/635/747) mounted in Ti cube polychroic, a Kinetix 22 back-illuminated sCMOS camera (Photometrics), and a stage-top incubator set to 37°C (Tokai Hit). ExRai AMPKAR and HyPer7 was imaged using a 380/20 and 475/20 LEDs coupled with a 390 nm or 475 nm excitation filter, respectively, and 515 nm emission filter. iATPSnFR2 was imged using 475/20 LEDs coupled with a 475 nm excitation filter and 515 nm emission filter. mScarlet and mCherry were imaged using a 575/25 LED and 575 nm excitation filter and 595 nm emission filter. Exposure times were set at 25-100 ms. Images were acquired every 30 s for up to 1 hr using NIS-Elements software (Nikon).

Image analysis was performed using published MATLAB code.^14,63^ For time-lapse experiments, regions of interest (ROIs) were drawn within randomly selected individual cells. For localized biosensors, ROIs were drawn around subcellular compartments, including mitochondria, lysosomal regions, and the nucleus. Raw fluorescence intensities were corrected for background fluorescence by subtracting signal from a cell-free area. For ExRai AMPKAR and HyPer7, excitation ratios (Ex480/Ex400) were calculated at each time point.^64^ For iATPSnFR2, the background-corrected intensity was calculated. All ratios or intensities were normalized to the baseline (pre-stimulation, typically 0-5 min) values. Maximum ratio changes (ΔR/R₀) were calculated using (R_max_-R_0_)/R_0_, where R_0_ is the ratio at the beginning of the experiment. For iATPSnFR, maximum or minimum fluorescence changes (F/F_0_) were calculated using (F_max_-F_0_/F_0_) or (F_min_-F_0_/F_0_), respectively. Time to half maximum and minimum calculations were performed using a custom R script to determine the time point at which each fluorescence trace reached half of its maximal (or minimal) response following stimulation. Graphing and statistical analysis were performed using GraphPad Prism 10 (GraphPad Software).

### Confocal imaging and image analysis

Confocal microscopy images and Z-stacks were acquired using a Leica STELLARIS 5 confocal microscope (Leica Microsystems) equipped with a DMOD WLL (440-790 nm), HC PL APO 63x/1.40 immersion CS2 lens, and a Power HyD S detector. LasX software (Leica) was used to control the microscope. Z-stacks were processed by generating maximum intensity projections in FIJI (ImageJ). Pseudo color images were also created using FIJI.

For assessing biosensor localization, cells expressing organelle-targeted ExRai AMPKAR were stained with either Invtrogen NucBlue Live ReadyProbes Reagent (Fisher Scientific Cat #R37605), MitoTracker Deep Red (Thermo Scientific Cat #M22426), or LysoTracker Deep Red (Thermo Scientific Cat #L12492) for 10 minutes. Merged, pseudo color images were created using FIJI.

### Western blotting

Cells were plated in 10 cm dishes in growth medium and grown to ∼90% confluency prior to treatment with small molecules. Following treatment, cells were washed with cold DPBS and lysed in lysis buffer (mammalian protein extraction reagent (M-PER, Thermo Fisher Cat #78501), Pierce™ protease inhibitor mini tablets, EDTA-free (Thermo Scientific Cat #A32955), Pierce™ phosphatase inhibitor mini tablets (Thermo Scientific Cat #A32957)). Lysates were vortexed, incubated on ice, and clarified by centrifugation at 12,000 x g for 20 min at 4°C. Protein concentration was determined via BCA assay (Thermo Scientific Cat #23225). Protein samples were separated using 4-15% SDS-PAGE (BioRad Cat #4568084) and transferred to nitrocellulose membranes (Bio-Rad Cat #1704158). Membranes were blocked in 1% BSA (Fisher Scientific Cat #52-171-00G) in TBST (20mM Tris, 150 mM NaCl, 0.05% Tween-20) and incubated with indicated primary antibodies overnight at 4°C. After washing, membranes were incubated with HRP-conjugated secondary antibodies for 1 hr at room temperature, and proteins were visualized using Clarity™ Western ECL substrate (BioRad Cat #1705060) and imaged using Azure imager 600.

### Proteomics sample preparation

MEFs were plated in 10 cm dishes in growth medium. Cells were treated with small molecules for indicated time. Cells were washed with ice cold PBS and then lysed in 8M urea lysis buffer (8M urea (Promega Cat #V3171), 100 mM tris (pH 8.0; Corning Cat #46-031-CM), 250 mM NaCl (Invitrogen Cat #AM9759), 50mM ammonium bicarbonate (Sigma-Aldrich Cat #09830-500G), cOmplete™ Mini Protease Inhibitor Cocktail (Roche Cat #11836153001) and PhosSTOP phosphatase inhibitor (Roche Cat #4906837001)) and stored on ice until sonication. Lysed samples were sonicated on ice using a probe sonicator 2x for 15 seconds at 10% amplitude, and protein was quantified via BCA assay. Proteins were reduced with 10 mM tris(2-carboxyethyl)phosphine (TCEP; Thermo Scientific Cat #20491) for 30 min at room temperature with shaking at 1500 rpm, followed by alkylation with 40 mM 2-chloroacetamide (Thermo Scientific Cat #A15238.30) for 30 min at room temperature with shaking at 1500 rpm. Samples were diluted 5-fold with 100 mM Tris-HCl (pH 8.0) to reduce urea concentration and digested overnight at room temperature with Lys-C (1:50, w/w) (Wako Cat #123175-82-6) and trypsin (1:100, w/w; Promega Cat #V5280) in a thermomixer at 800 rpm. Digestion was quenched by addition of 10% trifluoroacetic acid (TFA; Fisher Scientific Cat #A116-50) diluted in HPLC-grade water (Fisher Scientific Cat #AA47146M6) to a final pH of 2-3, insoluble material was pelleted by centrifugation, and peptides were desalted on Oasis HLB Vac Cartridges (30 mg sorbet per cartridge; Waters Cat #50-785-991) using standard activation (80% acetonitrile (ACN; Fisher Scientific Cat #A955-4)/0.1% TFA), equilibration wash (0.1% TFA), and elution (50% ACN/0.25% formic acid (FA; Fisher Scientific Cat #A117-50)). 10% of the elute was processed for global proteomics and the remaining 90% was processed for phosphopeptide enrichment.

For global proteomics, eluted peptides were dried by vacuum centrifugation and resuspended in 0.1% FA in water (Fisher Scientific Cat #LS118-4). For phosphoproteomics, Ti-IMAC HP beads (Resyn Biosciences Cat #MR-THP010) were used for phosphopeptide enrichment according to the manufacturer’s protocol with binding buffer (80% ACN, 5% TFA, 0.1 M glycolic acid), sequential washes (binding buffer, 60% ACN/1% TFA/200 mM NaCl, 60% ACN/1% TFA and HPLC-grade water), and elution with 1% ammonium hydroxide. The eluate was immediately neutralized with 10% FA, dried by vacuum centrifugation, and resuspended in 0.1% FA for LC-MS analysis.

### Mass spectrometry proteomics acquisition and analysis

Peptides were analyzed on a timsTOF HT mass spectrometer (Bruker Daltonics) coupled to a Vanquish Neo UHPLC system (Thermo Fisher). Samples were trapped on a PepMap Neo Trap Column (Thermo Scientific Cat #174500) and separated by reversed-phase chromatography on an Aurora Elite C18 column (15 cm length, 75 μm diameter, 1.7 μm particle size for captive spray; ESI Solutions Cat #AUR3-15075C18-CSI). The column was maintained at 50°C using a column oven for Bruker Captive Spray source (Sonation Lab Solutions).

Ionization was performed using a CaptiveSpray source (Bruker Daltonics) at 1700 V. Mobile phase A was 0.1% FA in water (Fisher Scientific Cat #LS118-1) and mobile phase B was 0.1% FA in ACN (Fisher Scientific Cat #LS120-1). The 45-minute LC gradient was run with a flow rate of 300 nL/min set as follows: 5-35% B over 37 min, then to 45% B over 4 min, then to 60% B over 1 min, and then to 95% B for 3 minutes. On the timsTOF HT, data were acquired in dia-PASEF mode. Equal-size windows of 22 Da were designed with an overlap of 1 Da to maximize the precursors ion coverage for further MS/MS. The ion accumulation time and ramp times in the dual TIMS analyzer were set to 100 ms each. In the ion mobility (1/K0) range 0.6 to 1.6 Vs cm-2, the collision energy was linearly decreased from 59 eV at 1/K0 = 1.43 Vs cm-2 to 20 eV at 1/K0 = 0.65 Vs cm-2 to collect the MS/MS spectra in the mass range 275.9 to 1368.9 Da. The estimated mean cycle time was 1.38 s.

Raw files generated from the MS were processed with Spectronaut (Biognosys v19.9.250512.62635) with the directDIA+ (Deep) search algorithm. Carbamidomethylation (cysteine) was set as a fixed modification for database search. Acetylation (protein N-term), oxidation (methionine), and phosphorylation (serine, threonine, tyrosine) were set as variable modifications. Reviewed *Mus musculus* protein sequences (downloaded from UniProt, February 23, 2024) were used for spectral matching.

The false discovery rates for the PSM, peptide, and protein groups were set to 0.01, and the minimum localization threshold for PTM was set to zero. For MS2-level area-based quantification, the cross-run normalization option was unchecked, and the probability cutoff was set to 0 for the PTM localization. Quantitative analysis was performed in R (v.4.4.1). Initial quality control analyses, including inter-run clustering, correlations, principal component analysis (PCA), peptide and protein counts, and intensities were completed using in-house code. MSstats^65^ was parameterized to perform normalization by median equalization, no imputation of missing values, and median smoothing (Tukey’s median polish) to combine intensities for multiple peptide ions or fragments into a single intensity for their protein group, and statistical tests of differences in intensity between conditions. Default settings for MSstats were used for adjusted P values. By default, MSstats uses the Student’s t-test for P value calculation. Statistical comparisons of phosphorylation changes were computed between WT MEFs treated with hydrogen peroxide or water, with or without AMPK inhibition by SBI-0206965.

Kinase activity analysis was performed on phosphoproteomics data to assess signaling changes downstream of oxidative stress-induced AMPK activity. Log2 fold changes in phosphosite intensities were calculated between treatment conditions, and enrichment of kinase activity was inferred using kinase-substrate annotations from the database, as done previously.^66,67^ For each kinase, activity was inferred by testing whether its substrates showed systematic shifts in Log2 fold change relative to the global distribution of phosphosite changes. The resulting Z-test statistics were converted to z-scores representing inferred kinase activity.^68^ *LC-MS metabolomics*

MEFs were plated in 6-well plates (Genesee Scientific Cat #25-105) in growth medium, and in triplicates of each condition. Cells were treated with hydrogen peroxide and/ or SBI-0206965 for indicated time. Packed cell volume (PCV) was determined using an additional well without treatment. The well was briefly washed and incubated with Trypsin (Thermo Scientific Cat #12604013), and pipetted into PCV tubes (Millipore Sigma Cat #Z760986) for centrifugation at 2,500 rcf for 1 min. The treated dishes were washed once with HBSS and 50 µL of 80% ethanol per 1 µL PCV was added, and the plate was incubated on dry ice for 10 min. The wells were then scraped with cell scrapers (Genesee Scientific Cat #25−270) and the mixture was transferred to microcentrifuge tubes for centrifugation at 16k rcf for 10 min at 4°C. The supernatant was then transferred to clean microcentrifuge tubes and stored at −80° until LC-MS analysis.

On the day of LC-MS analysis, samples were spun again at 16,000 rcf for 10 min at 4°C, and the supernatant was transferred into LC-MS tubes for analysis. A quadrupole-orbitrap mass spectrometer (Exploris 480, Thermo Fisher Scientific) operating in polarity-switching mode was coupled to hydrophilic interaction liquid chromatography (HILIC) via heated electrospray ionization. One full scan was performed from m/z 70−1000 at 90,000 resolution. Additional tSIM scans at 90,000 resolution with an isolation window of ±0.7 m/z were run in positive mode for potential malonyl-CoA ions (C_21_H_38_N_7_O_19_P_3_S) with AGC target set to standard and Maximum Injection Time Mode set to Auto. The inclusion list included singly charged ions: ^+^H adduct m/z 854.1229, ^+^NH4 adduct m/z 871.1494, ^+^K adduct m/z 892.0788, ^+^Na adduct m/z 876.1048, and no adduct m/z 853.1151. ddMS^2^ scans were also ran for these ions with stepped HCD Collision Energies of 10 and 40, Orbitrap resolution of 30,000, ACG target set to standard, and Maximum Injection Time Mode set to Auto. LC separation was achieved with a polymeric ZIC-pHILIC column (2.1 mm × 150 mm × 5 μm particle size, 100 Å pore size; Merck KGaA, Darmstadt, Germany) using a gradient of solvent A (5 mM NH_4_OAc, pH ∼9.8) and solvent B (95:5 acetonitrile/water). Flow rate was 0.17 mL/min. The LC gradient was: 0–0.5 min, 90% B; 0.5–15.0 min, linear decrease to 30% B; 15.0–20.0 min, hold at 30% B; 20.0–21.5 min, return to 90% B; 21.5–30.0 min, re-equilibrate at 90% B. Autosampler temperature was 4 °C, and the injection volume was 10 µL. Malonyl-CoA peak identification was verified by running a standard with malonyl-CoA lithium salt (Sigma-Aldrich Cat #M4263), which resulted in a detectable peak at m/z 854.1229 (RT 14.63) and MS² diagnostic fragments characteristic of malonyl-CoA: m/z 428.0, 347.1, 303.1, 136.1, and 99.1. Data were converted to mzXML format with msconvert, and ion counts were exported using MAVEN 2.49.

### RT-qPCR

Total RNA was extracted using the RNeasy Mini Kit (Qiagen Cat #74106) according to the manufacturer’s instructions, and RNA concentration and purity were assessed by Nanodrop. For each sample, gene expression was measured using the iTaq™ Universal SYBR® Green One-Step Kit (Bio-Rad Cat #1725150) in 10 µL reactions containing 5 µL of 2× SYBR One-Step Master Mix, 0.125 µL iScript Reverse Transcriptase, 1 µL of each forward and reverse primer (300 nM final), 1–2 µL total RNA (typically 500 ng), and nuclease-free water. Reactions were performed in a Bio-Rad CFX96 thermal cycler or Thermo-Fischer Quant Studio 5 using the following program: 10 min at 50 °C (RT), 1 min at 95 °C (enzyme activation), followed by 40 amplification cycles (10 s at 95 °C, 30 s at 60 °C), and a melt curve from 65 °C to 95 °C. All samples were run in technical triplicate. No-RT and no-template controls were included to verify RNA specificity and rule out contamination. Relative mRNA expression of *CAT*, *NQO1*, and *GPX1* were quantified by the ΔΔCt method^69^ (Ct values with confidence below 0.75 were discarded), normalized to *GAPDH*, and plotted as relative mRNA expression compared to untreated control samples.

### Immunostaining

Cells were seeded and grown to 70-80% confluency in 6-well glass bottom plates (Cellvis, Cat #P06-1.5H-N) prior to treatment with small molecules. Following treatment, cells were washed with cold 1x PBS (Thermo Fisher, Cat #10010023) three times for five minutes and then fixed with 4% (v/v) paraformaldehyde (Fisher Scientific, Cat #50-980-487) for 20 min at room temperature. Post-fixation, cells were washed three times for 5 minutes each with 1x PBS and then permeabilized in 1x PBS with 0.1% Triton X-100 (Thermo Fisher, Cat #A16046.AE) and 5% BSA (Tocris Biosciences, Cat #5217) per well. After blocking, NRF2 (D1Z9C) Rabbit Monoclonal Antibody (Cell Signaling Technology, Cat #12721S) was added to the wells at a 1:500 dilution and rocked overnight at 4°C. The next day, cells were washed three times for five minutes each with 1x PBS and then incubated and rocked with HRP-Goat anti-Rabbit IgG (H+L) Highly Cross-Adsorbed Secondary Antibody, Alexa Fluor Plus 488 (1:200; Thermo Scientific, Cat #A-32731), protected from light, for 1 hour at room temperature. Post-incubation, the cells were washed three times for five minutes each with 1x PBS, and NucBlue (Thermo Fisher, Cat #R37605) was added during the second wash. The dishes were imaged one day post using confocal microscopy.

Nuclear-to-cytoplasmic (N/C) intensity ratios were quantified using CellProfiler. Two-channel grayscale 16-bit TIFF images were used, where channel 00 contained the nuclear stain and channel 01 contained the signal of interest. Nuclei were identified from the nuclear channel using global Otsu thresholding with shape-based declumping to separate touching nuclei. Whole-cell boundaries were then defined by propagating outward from the identified nuclei using the rescaled signal channel image. Cells contacting the image border were excluded from analysis. Cytoplasmic regions were defined by subtracting the nuclear area from the whole-cell area. Mean fluorescence intensity of the signal channel was measured separately within nuclear and cytoplasmic compartments for each cell, and the nuclear-to-cytoplasmic intensity ratio was calculated from these values.

### Statistics and reproducibility

Figure preparation and statistical analysis were performed using GraphPad Prism 10 or R. For comparison of two data sets, nonparametric Mann-Whitney tests were used. For comparisons across more than two groups, a nonparametric Kruskal-Wallis test was performed, followed by multiple comparisons when appropriate. Cross correlation analysis was performed using the MATLAB xcorr function to quantify temporal relationships between signals, and correlation coefficients were plotted as a function of lag. Statistical significance was defined as p < 0.05 with a 95% confidence interval. The number of trials, number of independent experiments, technical replicates, and statistical tests used are reported in all figure legends. Where appropriate, data was plotted using the SuperPlots method.^70^ All dot plots shown depict the mean ± SEM.

## Data Availability Statement

All data are available within the main manuscript and the Supporting information.

Our phosphoproteomics dataset was uploaded to the Proteomics IDEntifications Database (PRIDE) at www.ebi.ac.uk/pride/ under project accession PXD070516. For reviewers, the token is “4GPe68Aq5gNG” and can be accessed with Username = and Password = ivy1kFMkTHHd. Metabolomics data will be shared on Metabolomics Workbench.

## Code Availability Statement

When applicable, custom code is available via Github (https://github.com/dlschmitt).

## Author Information

Corresponding Author

Danielle L. Schmitt- Department of Chemistry and Biochemistry, Molecular Biology Institute, Institute for Quantitative and Computational Biology, University of California Los Angeles, Los Angeles, California, 90095, United States of America

## Authors

Kasey Parks— Department of Chemistry and Biochemistry, University of California Los Angeles, Los Angeles, California, 90095, United States of America

Arnav Jhawar— Department of Chemistry and Biochemistry, University of California Los Angeles, Los Angeles, California, 90095, United States of America

Alexia Andrikopoulos— Department of Chemistry and Biochemistry, University of California Los Angeles, Los Angeles, California, 90095, United States of America

Declan Winters— Department of Human Genetics, David Geffen School of Medicine and Institute for Quantitative and Computational Biosciences, University of California Los Angeles, Los Angeles, California, 90095, United States of America

Teagan S. Dean— Department of Chemistry and Biochemistry, University of California Los Angeles, Los Angeles, California, 90095, United States of America

Edmund D. Kapelczak— Department of Molecular and Medical Pharmacology, David Geffen School of Medicine, University of California Los Angeles, Los Angeles, California, 90095, United States of America

Tara TeSlaa— Department of Molecular and Medical Pharmacology, David Geffen School of Medicine and Molecular Biology Institute, University of California Los Angeles, Los Angeles, California, 90095, United States of America

Mehdi Bouhaddou— Department of Microbiology, Immunology, and Molecular Genetics, Molecular Biology Institute, and Institute for Quantitative and Computational Biosciences, University of California Los Angeles, Los Angeles, California, 90095, United States of America

## Author Contributions

K.P. and D.L.S conceived the project. K.P. and A.J. performed imaging experiments. K.P., A.A., and D.W. performed proteomics. A.A., D.W., and M.B. analyzed proteomics data. A.J. and A.A. performed RT-PCR. A.A., A.J., and K.P. performed western blots. K.P., T.D.R., and E.D.K., performed metabolomics. K.P. and E.D.K. analyzed metabolomics data. D.L.S., T.T., and M.B. oversaw all experiments. K.P. and D.L.S wrote the manuscript. K.P. and A.A. prepared the figures. All authors contributed to the final version of the manuscript.

## Acknowledgements

We thank Reuben Shaw and David Shackelford for sharing of cell lines, Andrew Wojtovich for sharing mito-HyPer7, Katie Vineall for her assistance with confocal microscopy, Sandya Yadav for assistance with RT-qPCR, Peter Mullen and Steven Pilley for training in metabolomics data analysis and presentation, Jack Scully for assistance with molecular cloning, Alex Hoffmann and all members of the Schmitt Lab for their helpful discussion. This work was supported by the National Institutes of Health (DP2GM154012 to D.L.S., R35GM160071 to M.B., T32GM145388 to D.M.W., and T32GM007185 to A.A.), the UCLA Faculty Career Development Award (to D.L.S.), the UCLA Society of Hellman Fellows Award (to D.L.S.), UCLA Academic Senate Council on Research (to D.L.S.), the Chan-Zuckerberg Initiative (MET-000000000151 to T.T. and D.L.S.), and UCLA Department of Chemistry and Biochemistry Summer Undergraduate Research Fellowship (to K.P.). We acknowledge the UCLA AIDS Institute, the James B. Pendleton Charitable Trust, and the McCarthy Family Foundation for their generous support of our research.

## References

1. Hardie, D. G., Ross, F. A. & Hawley, S. A. AMPK: a nutrient and energy sensor that maintains energy homeostasis. Nat. Rev. Mol. Cell Biol. 13, 251–262 (2012).

2. Lin, S.-C. & Hardie, D. G. AMPK: Sensing Glucose as well as Cellular Energy Status. Cell Metab. 27, 299–313 (2018).

3. Davies, S. P. et al. Purification of the AMP-activated protein kinase on ATP-γ-Sepharose and analysis of its subunit structure. Eur. J. Biochem. 223, 351–357 (1994).

4. Dasgupta, B. & Chhipa, R. R. Evolving Lessons on the Complex Role of AMPK in Normal Physiology and Cancer. Trends Pharmacol. Sci. 37, 192–206 (2016).

5. Ross, F. A., Jensen, T. E. & Hardie, D. G. Differential regulation by AMP and ADP of AMPK complexes containing different γ subunit isoforms. Biochem. J. 473, 189–199 (2016).

6. Khan, A. S. & Frigo, D. E. A spatiotemporal hypothesis for the regulation, role, and targeting of AMPK in prostate cancer. Nat. Rev. Urol. 14, 164–180 (2017).

7. Shackelford, D. B. & Shaw, R. J. The LKB1–AMPK pathway: metabolism and growth control in tumour suppression. Nat. Rev. Cancer 9, 563–575 (2009).

8. Hawley, S. A. et al. Calmodulin-dependent protein kinase kinase-β is an alternative upstream kinase for AMP-activated protein kinase. Cell Metab. 2, 9–19 (2005).

9. Shaw, R. J. et al. The tumor suppressor LKB1 kinase directly activates AMP-activated kinase and regulates apoptosis in response to energy stress. Proc. Natl. Acad. Sci. U. S. A. 101, 3329–3335 (2004).

10. Anderson, K. A. et al. Hypothalamic CaMKK2 Contributes to the Regulation of Energy Balance. Cell Metab. 7, 377–388 (2008).

11. Garcia, D. & Shaw, R. J. AMPK: Mechanisms of Cellular Energy Sensing and Restoration of Metabolic Balance. Mol. Cell 66, 789–800 (2017).

12. Afinanisa, Q., Cho, M. K. & Seong, H.-A. AMPK Localization: A Key to Differential Energy Regulation. Int. J. Mol. Sci. 22, 10921 (2021).

13. Jhawar, A., Parks, K. & Schmitt, D. L. Illuminating compartmentalized AMPK signaling in single cells. Biochem. J. 483, 849–860 (2026).

14. Schmitt, D. L. et al. Spatial regulation of AMPK signaling revealed by a sensitive kinase activity reporter. Nat. Commun. 13, 3856 (2022).

15. Hinchy, E. C. et al. Mitochondria-derived ROS activate AMP-activated protein kinase (AMPK) indirectly. J. Biol. Chem. 293, 17208–17217 (2018).

16. Auciello, F. R., Ross, F. A., Ikematsu, N. & Hardie, D. G. Oxidative stress activates AMPK in cultured cells primarily by increasing cellular AMP and/or ADP. FEBS Lett. 588, 3361–3366 (2014).

17. Zmijewski, J. W. et al. Exposure to Hydrogen Peroxide Induces Oxidation and Activation of AMP-activated Protein Kinase*,. J. Biol. Chem. 285, 33154–33164 (2010).

18. Shao, D. et al. A redox-dependent mechanism for regulation of AMPK activation by Thioredoxin1 during energy starvation. Cell Metab. 19, 232–245 (2014).

19. Emerling, B. M. et al. Hypoxic activation of AMPK is dependent on mitochondrial ROS but independent of an increase in AMP/ATP ratio. Free Radic. Biol. Med. 46, 1386–1391 (2009).

20. Zhang, C.-S. et al. The lysosomal v-ATPase-Ragulator complex is a common activator for AMPK and mTORC1, acting as a switch between catabolism and anabolism. Cell Metab. 20, 526–540 (2014).

21. Zhang, H. et al. AMPK activation serves a critical role in mitochondria quality control via modulating mitophagy in the heart under chronic hypoxia. Int. J. Mol. Med. 41, 69–76 (2018).

22. Herzig, S. & Shaw, R. J. AMPK: guardian of metabolism and mitochondrial homeostasis. Nat. Rev. Mol. Cell Biol. 19, 121–135 (2018).

23. Drake, J. C., et al. Mitochondria-localized AMPK responds to local energetics and contributes to exercise and energetic stress-induced mitophagy. Proc. Natl. Acad. Sci. 118, e2025932118 (2021).

24. Pak, V. V. et al. Ultrasensitive Genetically Encoded Indicator for Hydrogen Peroxide Identifies Roles for the Oxidant in Cell Migration and Mitochondrial Function. Cell Metab. 31, 642–653.e6 (2020).

25. Carling, D., Sanders, M. J. & Woods, A. The regulation of AMP-activated protein kinase by upstream kinases. Int. J. Obes. 2005 32 Suppl 4, S55–59 (2008).

26. Goodwin, J. M. et al. An AMPK-independent signaling pathway downstream of the LKB1 tumor suppressor controls Snail1 and metastatic potential. Mol. Cell 55, 436–450 (2014).

27. iATPSnFR2: A high-dynamic-range fluorescent sensor for monitoring intracellular ATP | PNAS. https://www.pnas.org/doi/10.1073/pnas.2314604121.

28. Xiao, B. et al. Structure of mammalian AMPK and its regulation by ADP. Nature 472, 230–233 (2011).

29. Trelford, C. B. & Shepherd, T. G. Insights into targeting LKB1 in tumorigenesis. Genes Dis. 12, 101402 (2025).

30. Skoulidis, F. et al. Co-occurring genomic alterations define major subsets of KRAS-mutant lung adenocarcinoma with distinct biology, immune profiles, and therapeutic vulnerabilities. Cancer Discov. 5, 860–877 (2015).

31. Kottakis, F. et al. LKB1 loss links serine metabolism to DNA methylation and tumorigenesis. Nature 539, 390–395 (2016).

32. Faubert, B. et al. Loss of the tumor suppressor LKB1 promotes metabolic reprogramming of cancer cells via HIF-1α. Proc. Natl. Acad. Sci. 111, 2554–2559 (2014).

33. Hardie, D. G. AMP-activated protein kinase — a journey from 1 to 100 downstream targets. Biochem. J. 479, 2327–2343 (2022).

34. Egan, D. F. et al. Phosphorylation of ULK1 (hATG1) by AMP-Activated Protein Kinase Connects Energy Sensing to Mitophagy. Science 331, 456–461 (2011).

35. Kim, J., Kundu, M., Viollet, B. & Guan, K.-L. AMPK and mTOR regulate autophagy through direct phosphorylation of Ulk1. Nat. Cell Biol. 13, 132–141 (2011).

36. Anderson, K. A. et al. Hypothalamic CaMKK2 Contributes to the Regulation of Energy Balance. Cell Metab. 7, 377–388 (2008).

37. Tojkander, S., Ciuba, K. & Lappalainen, P. CaMKK2 Regulates Mechanosensitive Assembly of Contractile Actin Stress Fibers. Cell Rep. 24, 11–19 (2018).

38. Tamás, P. et al. Regulation of the energy sensor AMP-activated protein kinase by antigen receptor and Ca2+ in T lymphocytes. J. Exp. Med. 203, 1665–1670 (2006).

39. Dite, T. A. et al. AMP-activated protein kinase selectively inhibited by the type II inhibitor SBI-0206965. J. Biol. Chem. 293, 8874–8885 (2018).

40. Nostrand, J. L. V. et al. AMPK regulation of Raptor and TSC2 mediate metformin effects on transcriptional control of anabolism and inflammation. Genes Dev. 10.1101/gad.339895.120 (2020) doi:10.1101/gad.339895.120.

41. Matzinger, M., Fischhuber, K., Pölöske, D., Mechtler, K. & Heiss, E. H. AMPK leads to phosphorylation of the transcription factor Nrf2, tuning transactivation of selected target genes. Redox Biol. 29, 101393 (2020).

42. Hardie, D. G. AMP-activated protein kinase - a journey from 1 to 100 downstream targets. Biochem. J. 479, 2327–2343 (2022).

43. Bhattacharya, S. et al. STAT3 suppresses the AMPKα/ULK1-dependent induction of autophagy in glioblastoma cells. J. Cell. Mol. Med. 26, 3873–3890 (2022).

44. Chen, L. et al. NADPH production by the oxidative pentose-phosphate pathway supports folate metabolism. Nat. Metab. 1, 404–415 (2019).

45. TeSlaa, T., Ralser, M., Fan, J. & Rabinowitz, J. D. The pentose phosphate pathway in health and disease. Nat. Metab. 5, 1275–1289 (2023).

46. Kim, D. et al. Mitochondrial NADPH fuels mitochondrial fatty acid synthesis and lipoylation to power oxidative metabolism. Nat. Cell Biol. 27, 790–800 (2025).

47. Owen, J. B. & Butterfield, D. A. Measurement of oxidized/reduced glutathione ratio. Methods Mol. Biol. Clifton NJ 648, 269–277 (2010).

48. Joo, M. S. et al. AMPK Facilitates Nuclear Accumulation of Nrf2 by Phosphorylating at Serine 550. Mol. Cell. Biol. 36, 1931–1942 (2016).

49. Liang, J. et al. Baicalin Attenuates H2O2-Induced Oxidative Stress by Regulating the AMPK/Nrf2 Signaling Pathway in IPEC-J2 Cells. Int. J. Mol. Sci. 24, 9435 (2023).

50. Lee, H. M. et al. Concurrent loss of LKB1 and KEAP1 enhances SHMT-mediated antioxidant defence in KRAS-mutant lung cancer. Nat. Metab. 6, 1310–1328 (2024).

51. Nova, Z., Skovierova, H., Strnadel, J., Halasova, E. & Calkovska, A. Short-Term versus Long-Term Culture of A549 Cells for Evaluating the Effects of Lipopolysaccharide on Oxidative Stress, Surfactant Proteins and Cathelicidin LL-37. Int. J. Mol. Sci. 21, 1148 (2020).

52. Gonet-Surówka, A. & Dynarowicz-Latka, P. Cisplatin-resistant A549 non-small cell lung cancer cells sensitivity to 7-ketocholesterol. J. Steroid Biochem. Mol. Biol. 260, 107006 (2026).

53. Bellezza, I., Giambanco, I., Minelli, A. & Donato, R. Nrf2-Keap1 signaling in oxidative and reductive stress. Biochim. Biophys. Acta BBA - Mol. Cell Res. 1865, 721–733 (2018).

54. Li, T. & Zhu, H. LKB1 and cancer: The dual role of metabolic regulation. Biomed. Pharmacother. 132, 110872 (2020).

55. Zulato, E. et al. LKB1 loss is associated with glutathione deficiency under oxidative stress and sensitivity of cancer cells to cytotoxic drugs and γ-irradiation. Biochem. Pharmacol. 156, 479–490 (2018).

56. Kazgan, N., Williams, T., Forsberg, L. J. & Brenman, J. E. Identification of a nuclear export signal in the catalytic subunit of AMP-activated protein kinase. Mol. Biol. Cell 21, 3433–3442 (2010).

57. Lawrence, R. E. & Zoncu, R. The lysosome as a cellular centre for signalling, metabolism and quality control. Nat. Cell Biol. 21, 133–142 (2019).

58. Kiraly, S., Stanley, J. & Eden, E. R. Lysosome-Mitochondrial Crosstalk in Cellular Stress and Disease. Antioxid. Basel Switz. 14, 125 (2025).

59. Weinberg, F. et al. Mitochondrial metabolism and ROS generation are essential for Kras-mediated tumorigenicity. Proc. Natl. Acad. Sci. 107, 8788–8793 (2010).

60. Sanchez-Cespedes, M. et al. Tolerance and Resistance to Targeted Therapy in NSCLC: Emerging Concepts and Strategies. JTO Clin. Res. Rep. 7, 100944 (2025).

61. Cai, H., Zhang, F., Xu, F. & Yang, C. Metabolic reprogramming and therapeutic targeting in non-small cell lung cancer: emerging insights beyond the Warburg effect. Front. Oncol. 15, (2025).

62. Liu, H. & Naismith, J. H. An efficient one-step site-directed deletion, insertion, single and multiple-site plasmid mutagenesis protocol. BMC Biotechnol. 8, 91 (2008).

63. Frei, M. S. et al. Far-red chemigenetic kinase biosensors enable multiplexed and super-resolved imaging of signaling networks. Nat. Biotechnol. 1–10 (2025) doi:10.1038/s41587-025-02642-8.

64. Schmitt, D. L. Imaging Subcellular AMPK Activity Using an Excitation-Ratiometric AMPK Activity Reporter. Curr. Protoc. 3, e771 (2023).

65. Choi, M. et al. MSstats: an R package for statistical analysis of quantitative mass spectrometry-based proteomic experiments. Bioinformatics 30, 2524–2526 (2014).

66. Türei, D. et al. Integrated intra- and intercellular signaling knowledge for multicellular omics analysis. Mol. Syst. Biol. 17, e9923 (2021).

67. Bouhaddou, M. et al. The Global Phosphorylation Landscape of SARS-CoV-2 Infection. Cell 182, 685–712.e19 (2020).

68. Müller-Dott, S. et al. Comprehensive evaluation of phosphoproteomic-based kinase activity inference. Nat. Commun. 16, 4771 (2025).

69. Livak, K. J. & Schmittgen, T. D. Analysis of relative gene expression data using real-time quantitative PCR and the 2(-Delta Delta C(T)) Method. Methods San Diego Calif 25, 402–408 (2001).

70. Lord, S. J., Velle, K. B., Mullins, R. D. & Fritz-Laylin, L. K. SuperPlots: Communicating reproducibility and variability in cell biology. J. Cell Biol. 219, e202001064 (2020).

