## Supplemental Information for "LKB1 Spatially Regulates AMPK Activity Coordinate Cellular Oxidative Stress Response"

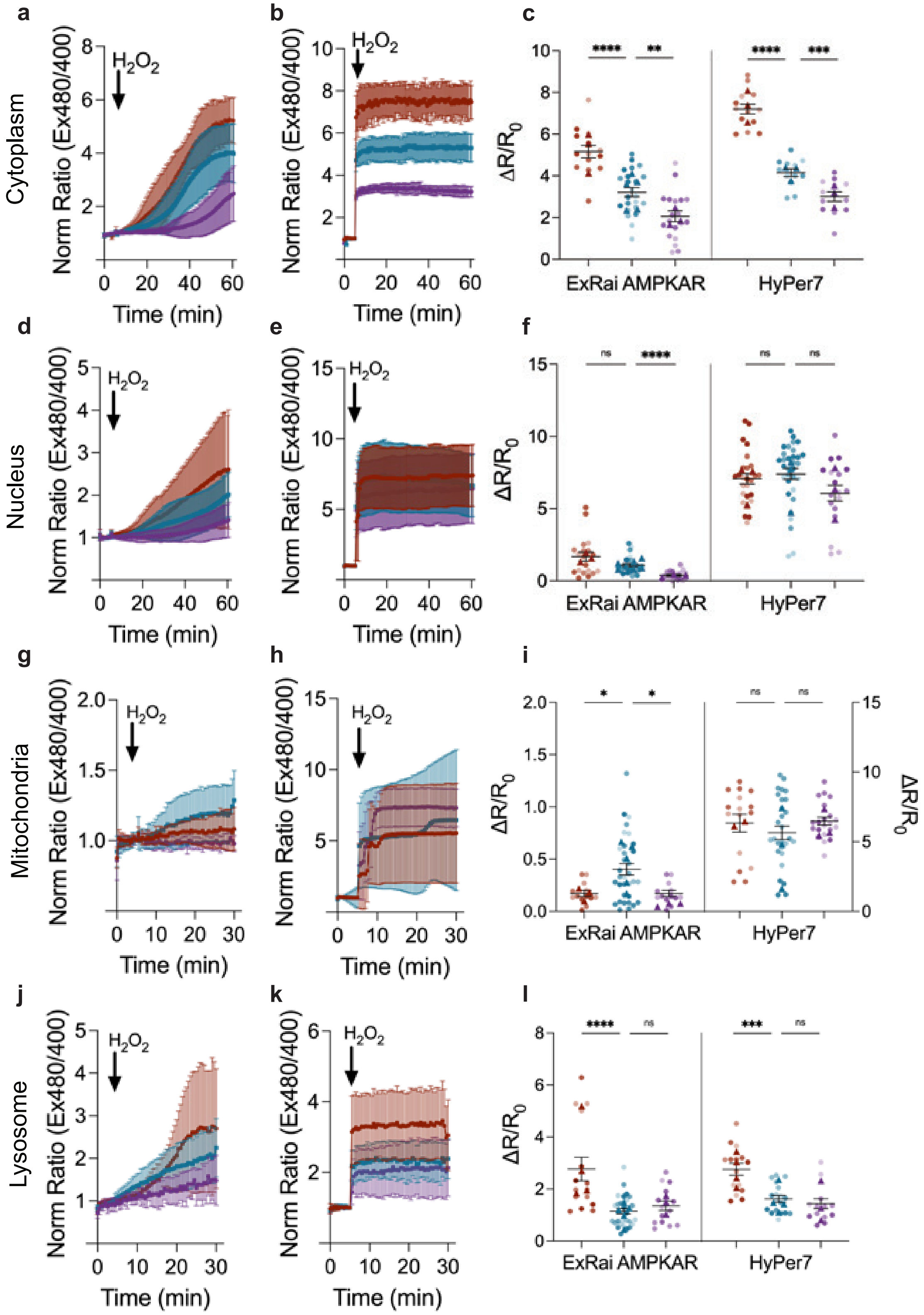


**Supplemental Figure 1. Spatial AMPK activity is sensitive to the cellular oxidation state.**

**a** Average response of ExRai AMPKAR in WT MEFs treated with 75 μM (purple, n=19 cells from three experiments), 150 μM (blue, reproduced from Fig. 1b), and 300 μM (red, n=14 cells from three experiments) $H_2O_2$.

**b** Average response of HyPer7 in WT MEFs treated with 75 μM (purple, n=13 cells from three experiments), 150 μM (blue, n=13 cells from three experiments), and 300 μM (red, n=14 cells from three experiments) $H_2O_2$.

**c** Maximum ratio changes of ExRai AMPKAR and HyPer7 in WT MEFs treated with varying concentrations of $H_2O_2$ (***p*=0.0042, ****p*=0.0005, ****p<0.0001 unpaired t-test, two-tailed

**d** Average response of NLS-ExRai AMPKAR in WT MEFs treated with 75 μM (purple, n=18 cells from three experiments), 150 μM (blue, reproduced from Fig. 1c), and 300 μM (red, n=18 cells from three experiments) $H_2O_2$.

**e** Average response of NLS-HyPer7 in WT MEFs treated with 75 μM (purple, n=18 cells from three experiments), 150 μM (blue, n=32 cells from three experiments), and 300 μM (red, n=26 cells from three experiments) $H_2O_2$.

**f** Maximum ratio changes of NLS-ExRai AMPKAR and NLS-HyPer7 in WT MEFs treated with varying concentrations of $H_2O_2$ (ns≥0.2408, *****p*<0.0001, unpaired *t*-test, two-tailed).

**g** Average response of mito-ExRai AMPKAR in WT MEFs treated with 75 μM (purple, n=15 cells from three experiments), 150 μM (blue, reproduced from Fig. 1d), and 300 μM (red, n=13 cells from three experiments) $H_2O_2$.

**h** Average response of mito-HyPer7 in WT MEFs treated with 75 μM (purple, n=20 cells from three experiments), 150 μM (blue, n=29 cells from three experiments), and 300 μM (red, n=22 cells from three experiments) $H_2O_2$.

**i** Maximum ratio change of mito-ExRai AMPKAR and mito-HyPer7 in WT MEFs treated with varying concentrations of $H_2O_2$ (ns≥0.02304, **p*=0.0191, *****p*<0.0001, unpaired *t*-test, two-tailed).

**j** Average response of lyso-ExRai AMPKAR in WT MEFs treated with 75 μM (purple, n=15 cells from three experiments), 150 μM (blue, reproduced from Fig. 1e), and 300 μM (red, n=15 cells from four experiments) $H_2O_2$.

**k** Average response of lyso-HyPer7 in WT MEFs treated with 75 μM (purple, n=15 cells from three experiments), 150 μM (blue, n=18 cells from four experiments), and 300 μM (red, n=15 cells from three experiments) $H_2O_2$.

**l** Maximum ratio changes of lyso-ExRai AMPKAR and lyso-HyPer7 in WT MEFs treated with varying concentrations of $H_2O_2$ (ns≥0.2963, ***p=0.0002, ****p<0.0001, unpaired *t*-test, two-tailed).

For all figures, time courses show the mean ± SD, dot plots show the mean ± SEM.

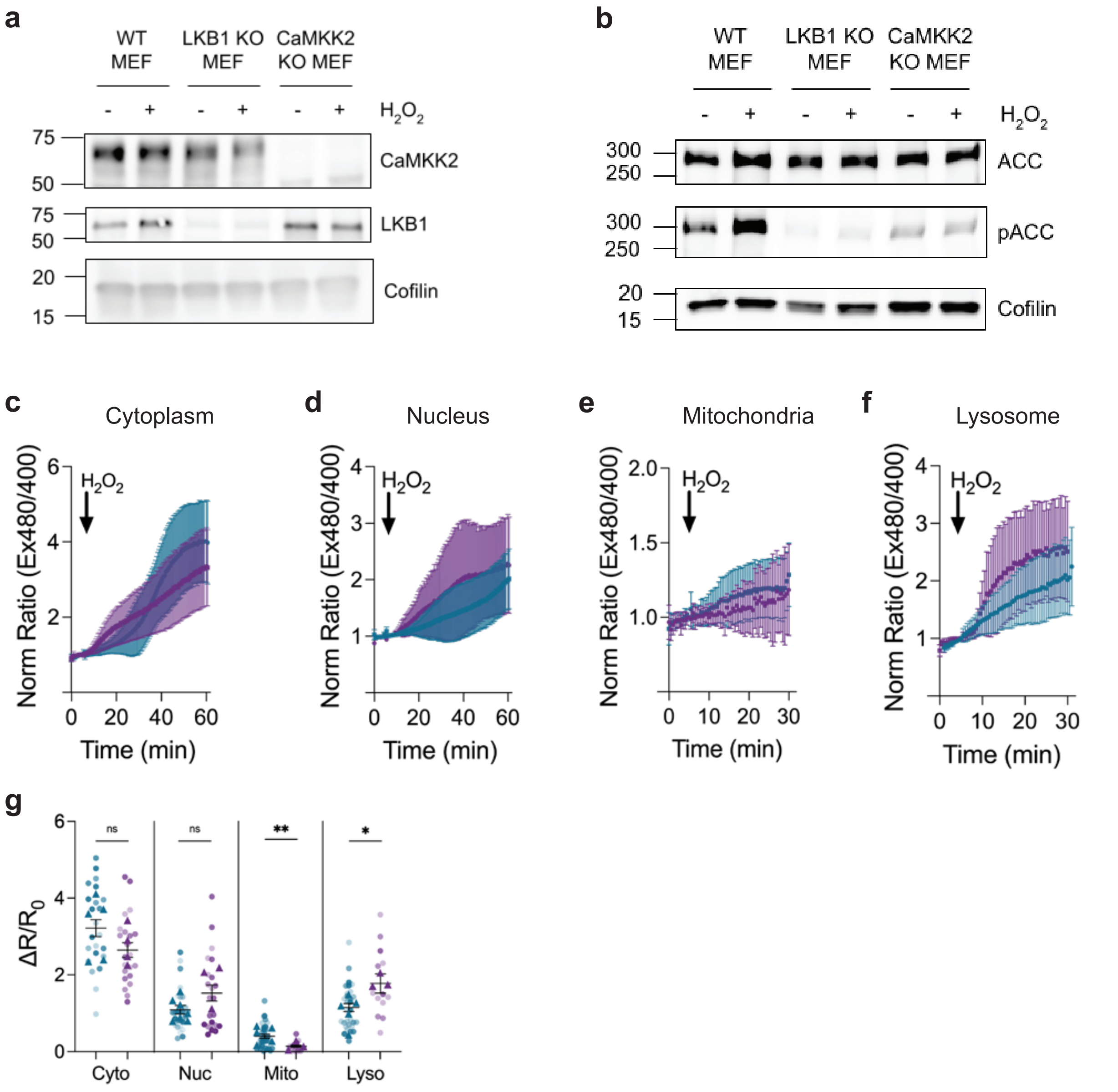


**Supplemental Figure 2. Impact of upstream kinases on oxidative stress-induced AMPK activity.**

**a** Representative western blot of WT, LKB1 KO, and CaMKK2 KO MEFs for CaMKK2 and LKB1.Full blots are shown in Source Data.

**b** Representative western blot of WT, LKB1 KO, and CaMKK2 KO MEFs treated with $H_2O_2$ (150 μM) or vehicle control and blotted for total and phospho-ACC. Full blots are shown in Source Data.

**c** Average response of ExRai AMPKAR in WT (blue, replotted from Fig 1b) and CaMKK2 KO (purple, n=25 from five experiments) MEFs treated with $H_2O_2$ (150 μM).

**d** Average response of NLS-ExRai AMPKAR in WT (blue, replotted from 1c) and CaMKK2 KO (purple, n=22 from three experiments) MEFs treated with $H_2O_2$ (150 μM).

**e** Average response of lyso-ExRai AMPKAR in WT (blue, replotted from 1e) and CaMKK2 KO (purple, n=13 from three experiments) MEFs treated with $H_2O_2$ (150 μM).

**f** Average response of mito-ExRai AMPKAR in WT (blue, replotted from 1d) and CaMKK2 KO (purple, n=23 from three experiments) MEFs treated with $H_2O_2$ (150 μM).

**g** Maximum ratio change ($\Delta R/R_0$) of organelle targeted ExRai AMPKAR in WT (blue) and CaMKK2 KO (purple) MEFs treated with $H_2O_2$ (150 μM), (ns>0.0357, ***p*=0.0020, unpaired t-test, two-tailed).

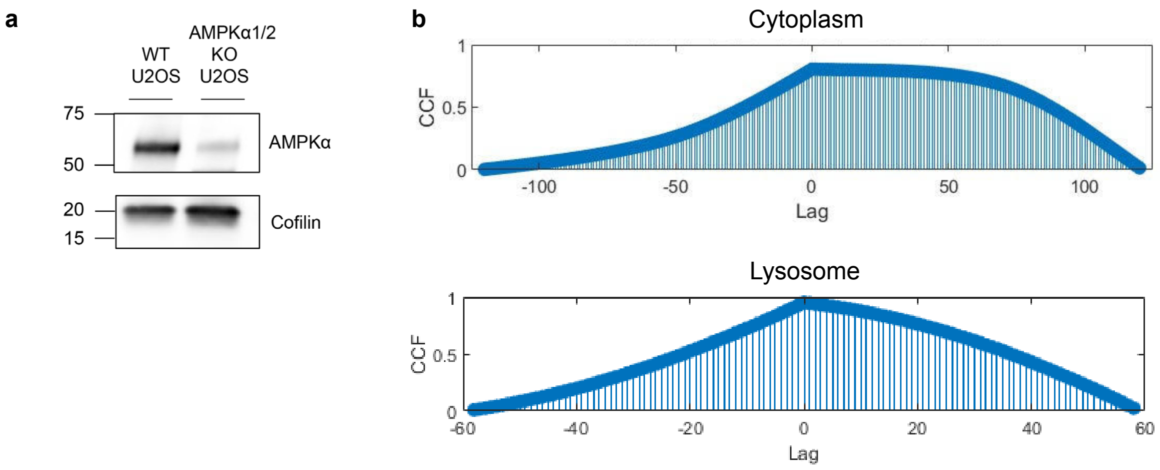


**Supplemental Figure 3. Phosphorylation of AMPK is required for activity under oxidative stress.**

**a** Representative western blot for U2OS and AMPKα1/2 KO U2OS for AMPKα. Full blots are shown in Source Data.

**b** Cross correlation of iATPSnFR2 between WT (signal 1) and LKB1 KO (signal 2) MEFs for the cytoplasm (top) and lysosome (bottom). Lag at time peaking at 0 indicates data are highly synchronized. Error bars represent SEM.

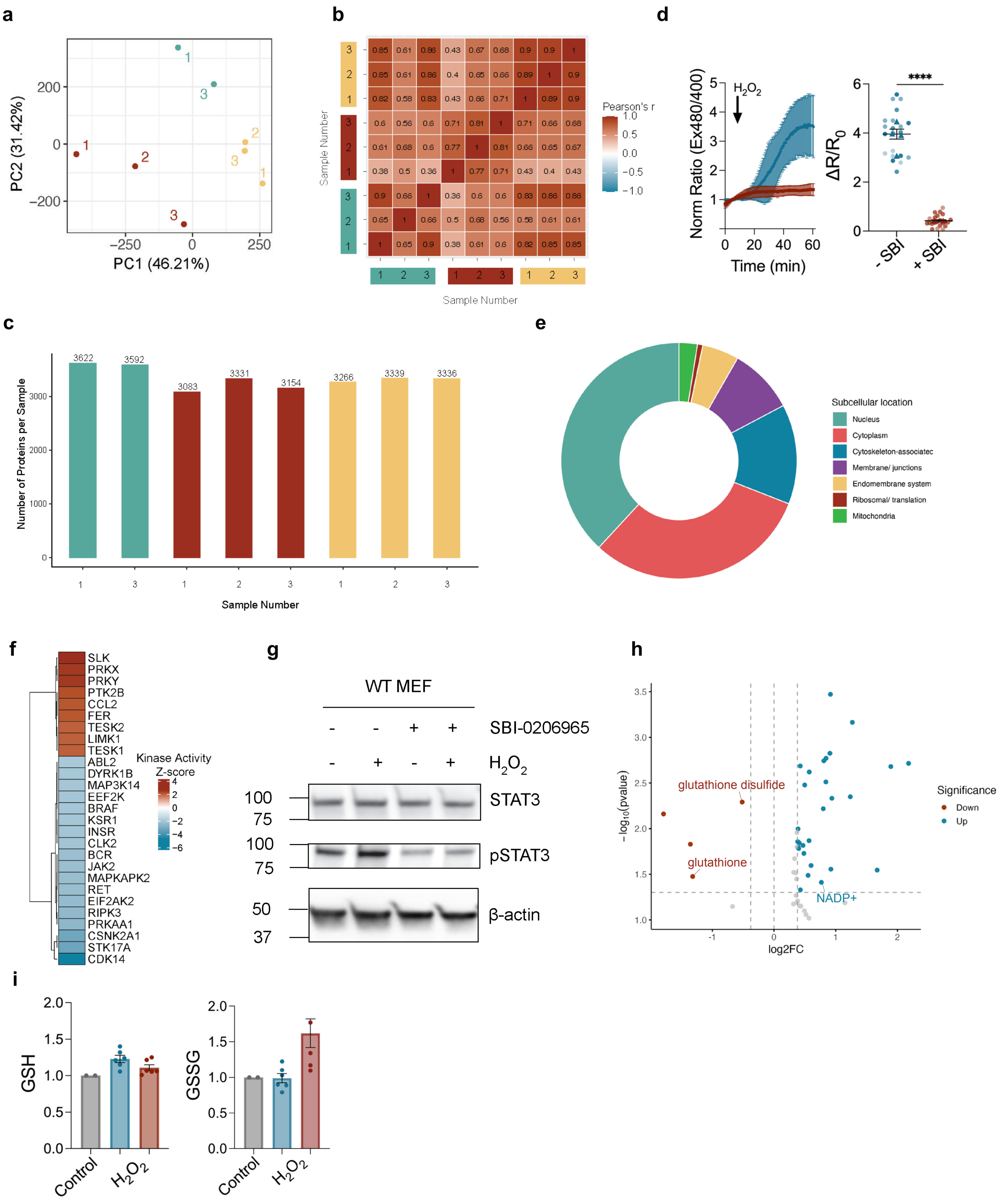


**Supplemental Figure 4. Phosphoproteomics and metabolomics analysis of AMPK-mediated signaling networks under oxidative stress.**

**a** Principal component analysis (PCA) of phosphoproteomics data from WT MEFs treated with $H_2O_2$ (150 μM) (blue), pre-treated with SBI-0206965 (red), and vehicle control (yellow). PC1 accounts for 46.21% and PC2 for 31.42% of variance.

**b** Pearson's correlation coefficient of phosphoproteomics data from WT MEFs treated with $H_2O_2$ (150 μM) (blue), pre-treated with 10 μM SBI-0206965 (red), and vehicle control (yellow). Sample number indicates technical replicate. Sample 2 from the $H_2O_2$ treated experimental condition was discarded due to pearson's correlation coefficient < 0.75.

**c** Total protein count of phosphoproteomics data from WT MEFs treated with $H_2O_2$ (150 μM) (blue), pre-treated with 10 μM SBI-0206965 (red), and vehicle control (yellow). Sample number indicates technical replicate.

**d** Average response of ExRai AMPKAR in WT MEFs (blue, n=23 cells from five experiments) and WT MEFs pre-treated with 10 μM SBI-0206965 (red, n=29 cells from three experiments) treated with $H_2O_2$ (150μM), along with maximum ratio change (****$p$<0.0001, unpaired $t$-test, two-tailed).

**e** Distribution of phosphopeptides by subcellular location from proteomics analysis.

**f** Heatmap showing the kinase activities of phosphorylated substrates from global MS phosphoproteomics analysis of WT MEFs treated with $H_2O_2$ (150 μM) measured by z-score, $p\leq$ 0.05. Red represents upregulated and blue represents downregulated in SBI-0206965 treated conditions.

**g** Representative western blot of WT MEFs treated with $H_2O_2$ (150 μM) with and without SBI-0206965 (10 μM) for total and phospho-STAT3. Full blots are shown in Source Data.

**h** Volcano plot highlighting changes in metabolites between WT MEFs treated with $H_2O_2$ (150 μM), with and without SBI-0206965 (10 μM). Colored dots depict downregulated (red, n= 4 metabolites) and upregulated (blue, n= 27 metabolites) metabolites in SBI-0206965 treated conditions at |FC > 1.3| and $p$<0.05 cutoffs.

**i** Ion count for GSH and GSSG in WT MEFs after treatment with $H_2O_2$ (150 μM) (blue, n=2 independent experiments with 2-3 technical replicates), and with pre-treatment with 10 μM SBI-0206965 (dark red, n=2 experiments with 2-3 technical replicates), normalized to vehicle control (grey, n=2 independent experiments with 1 technical replicate), (*$p\leq$0.0227, Kruskal-Wallis test).

### Supplementary Table 1. Primers used for molecular cloning and RT-qPCR

| Primer Number | Purpose | Sequence (5' to 3') |
|---|---|---|
| 1 | AMPKα2 T172A, forward | CCACAGCTCGCTCTTAAAATTCACCATCTGACATCATGTTG |
| 2 | AMPKα2 T172A, reverse | TTAAGAGCGAGCTGTGGCTCGCCCAATTATGCTGC |
| 3 | NQO1 (NM_008706), forward | GAGAAGAGCCCTGATTGTACTG |
| 4 | NQO1 (NM_008706), reverse | ACCTCCCATCCTCTCTTCTT |
| 5 | GPX (GID 14775), forward | CTCACCCGCTCTTTACCTTCCT |
| 6 | GPX (GID 14775), reverse | ACACCGGAGACCAAATGATGTACT |
| 7 | Catalase (NM_009804.2), forward | CAGATGGAGAGGCAGTCTATTG |
| 8 | Catalase (NM_009804.2), reverse | AAAGATCTCGGAGGCCATAATC |
| 9 | GAPDH (NM_008084), forward | GGTGTGAACCATGAGAAGTATGA |
| 10 | GAPDH (NM_008084), reverse | GAGTCCTTCCACGATACCAAAG |
